# Deep learning can predict multi-omic biomarkers from routine pathology images: A systematic large-scale study

**DOI:** 10.1101/2022.01.21.477189

**Authors:** Salim Arslan, Debapriya Mehrotra, Julian Schmidt, Andre Geraldes, Shikha Singhal, Julius Hense, Xiusi Li, Cher Bass, Jakob Nikolas Kather, Pahini Pandya, Pandu Raharja-Liu

## Abstract

We assessed the pan-cancer predictability of multi-omic biomarkers from haematoxylin and eosin (H&E)-stained whole slide images (WSI) using deep learning (DL) throughout a systematic study. A total of 13,443 DL models predicting 4,481 multi-omic biomarkers across 32 cancer types were trained and validated. The investigated biomarkers included a broad range of genetic, transcriptomic, proteomic, and metabolic alterations, as well as established markers relevant for prognosis, molecular subtypes and clinical outcomes. Overall, we found that DL can predict multi-omic biomarkers directly from routine histology images across solid cancer types, with 50% of the models performing at an area under the curve (AUC) of more than 0.633 (with 25% of the models having an AUC larger than 0.711). A wide range of biomarkers were detectable from routine histology images across all investigated cancer types, with a mean AUC of at least 0.62 in almost all malignancies. Strikingly, we observed that biomarker predictability was mostly consistent and not dependent on sample size and class ratio, suggesting a degree of true predictability inherent in histomorphology. Together, the results of our study show the potential of DL to predict a multitude of biomarkers across the omics spectrum using only routine slides. This paves the way for accelerating diagnosis and developing more precise treatments for cancer patients.

## Introduction

Studying the alterations at different levels of the molecular landscape helps better understand oncogenesis and cancer progression^1, 2^. In-depth analysis of the associations between the molecular aberrations and the tumour microenvironment has enabled the development of targeted therapies in various cancer types^3–5^. Genetic profiling has become an important tool^6^, especially for individuals who possess a higher risk of developing cancer due to genetic factors^7^. There is also an increasing need for alternative solutions to standard molecular and genomic profiling methods, which are often accountable for laboratory delays in the routine clinical workflow, as they take time to prepare, process, and analyse^8, 9^. In addition, expensive tests may not be routinely accessible to all patients^10^.

Meanwhile, there has been accumulating evidence suggesting that diagnostic histology images stained with hematoxylin and eosin (H&E) may contain information that can be used to infer molecular profiles directly from histological slides^11, 12^. Deep learning (DL) can effectively reveal differences in morphological phenotypes in malignancies, which in turn, enables the prediction of molecular profiles directly from H&E-stained whole slide images (WSIs)^11–15^. DL-based methods have been used to infer molecular alterations in various cancer types, including breast^9, 16–18^, colorectal^13, 15, 19, 20^, lung^12, 21^, gastric^22^, prostate^23^, skin^24^, and thyroid^25^ (see Echle et al. for an extensive review of DL applications for biomarker profiling from histology images^26^). More recently, pan-cancer studies have explored the links between genetic/molecular alterations and histomorphological features in H&E images. These studies showed that in almost all malignancies, DL methods can be used to infer a plethora of biomarkers directly from routine histology ^11, 14, 15, 27–29^.

Expanding upon previous work, we conducted a large-scale study to assess the feasibility of biomarker profiling from routine diagnostic slides with DL. The investigated biomarkers ranged from genomic, transcriptomic, proteomic, and metabolic alterations to various clinically-relevant downstream biomarkers (e.g. standard of care features, molecular subtypes, gene expressions, clinical outcomes, and response to treatment). We systematically evaluated predictability across all solid cancers studied by the Cancer Genome Atlas (TCGA) program. Our DL approach utilises an autoencoder network as a feature extractor that enables learning representations of the histology images which are relevant for the profiling task at hand. In contrast to previous studies, we have extended the scope of assessing pan-cancer predictability with DL to a wider range of cancer types (n=32) and to thousands of biomarkers across the central dogma of molecular biology (n=4,481). The systematic predictability of many of these biomarkers from routine histology has not been assessed at large scale before, including metabolic pathways and certain phenotypic or clinical outcomes such as drug responses.

Overall, we found that multi-omic biomarkers can be predicted directly from histo-morphology. Profiling mutations from histology was mostly feasible for the majority of genes tested and frequently mutated genes like *TP53* were predictable across multiple cancer types. While it was possible to predict the under-/over-expression status of transcriptomes and proteins to a certain degree, metabolic biomarkers were somewhat less predictable. We observed that the morphological visual characteristics could be detected with DL, enabling the prediction of molecular subtypes and well-established clinical biomarkers directly from WSIs. Repeating our experiments with an external dataset based on certain cancer types and biomarkers that have equivalents in the TCGA dataset, we obtained similar results, further confirming the general feasibility of predicting pan-cancer biomarkers from H&E slides. Considering various factors that may have an impact on predictability, such as sample size and prevalence of biomarker status, we conclude that there exists a degree of true predictability that may be associated with histomorphology.

## Results

### DL for molecular profiling from routine histology images

A convolutional neural network (CNN) was used for predicting molecular profiles from H&E images as illustrated in **Figure 1a** and explained in **Online Methods**. Our CNN consisted of an encoder (i.e. feature extractor), a decoder and a profiling module for classification. In conventional neural networks, morphological features acquired from histology images are directly correlated with the target molecular profile. In our approach, the combined classification and encoder-decoder architecture enable learning a better representation (i.e. encoding) of morphological images that are free of irrelevant features like image noise. The investigated biomarkers in our study included genetic alterations in driver genes, over-and under-expression of driver genes and relevant proteins, established biomarkers that are routinely used in clinical management, clinical outcomes such as overall survival and treatment responses, biomarkers that are highly relevant for prognosis and targeted therapies, including molecular subtypes and gene expression signatures (**Online Methods: Biomarker acquisition**). We assessed the predictive performance of each biomarker in a 3-fold cross-validation setting, where the cases with a valid biomarker status in each cohort were split into three random partitions (folds), each having similar proportions of positive/negative samples. We trained and tested three models per biomarker, each time keeping aside a different fold for testing and using the remaining ones for training. This setting ensured that a test prediction could be acquired for each patient and allowed us to assess the variance in biomarker performance. The predictability of a biomarker was measured with the area under the receiver operating characteristic curve (AUC). For each biomarker, we reported the performance as the average AUC across the three models (unless otherwise specified), accompanied with the standard deviation (denoted with ± where appropriate) measuring intra-marker variability. In total, 13,443 models were trained across 4,481 distinct biomarkers and 32 cancer types, with the following breakdown (**Figure 1b**): 1,950 single nucleotide variant (SNV) mutations in driver genes, 1,030 transcriptome expression level markers, 576 protein expression level markers, 450 metabolic pathways, 270 gene signature and molecular subtype markers, 160 markers related to clinical outcomes and treatment responses and 45 standard-of-care markers.

**Figure 1.**
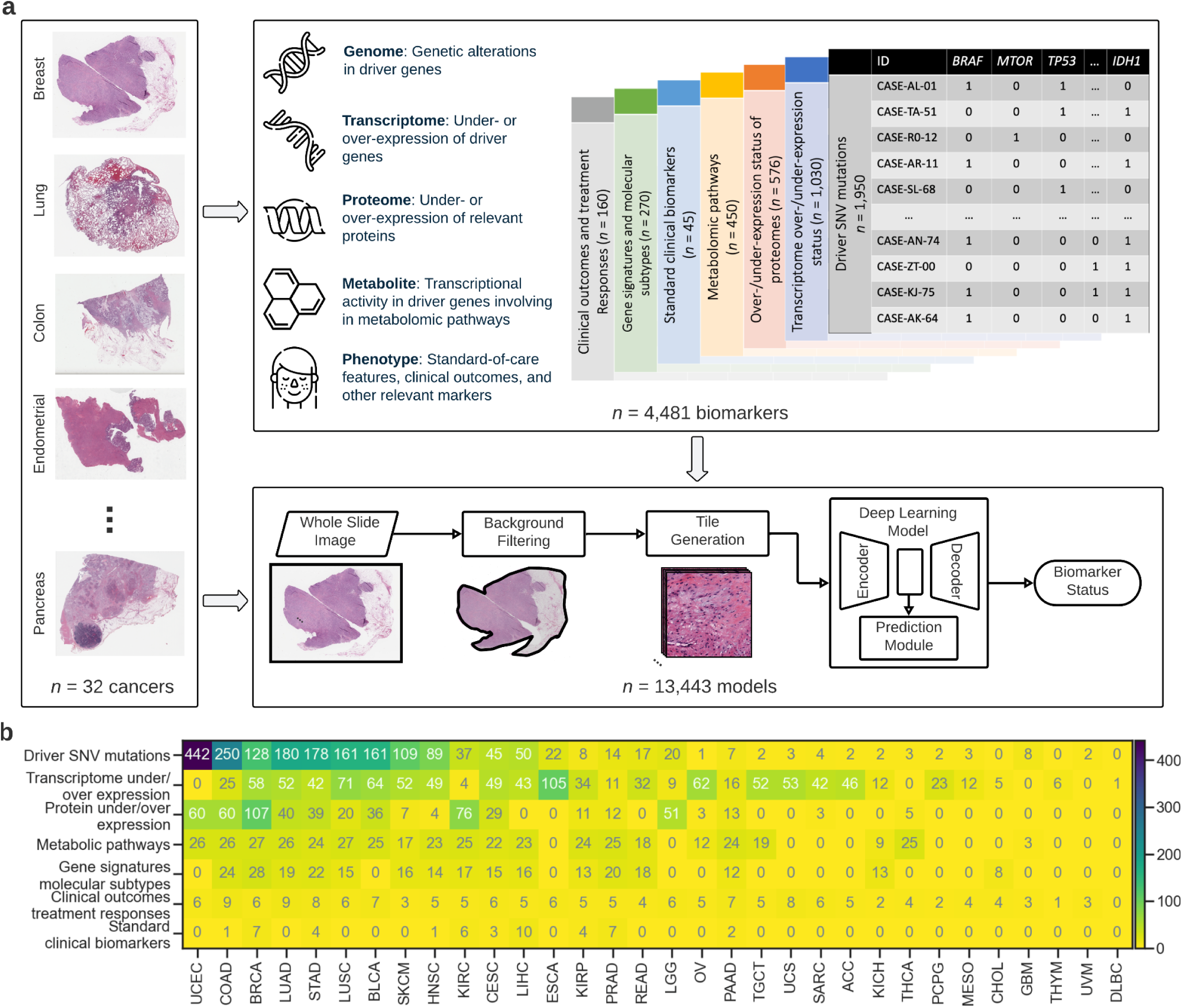
| Deep learning for molecular profiling from routine histology images. **(a)**: Overview of the pre-processing and training pipeline used for assessing the feasibility of predicting a plethora of biomarkers from genomic, transcriptomic, proteomic, metabolic to various clinically-relevant biomarkers (e.g. standard of care features, molecular subtypes, clinical outcomes, response to treatment) using H&E-stained whole slide images and deep learning. A CNN consisting of an encoder (i.e. feature extractor), a decoder and a classification module was used for predicting molecular profiles from H&E images (see **Online Methods: Pre-processing pipeline and training details**). A single CNN model was end-to-end trained from scratch for each biomarker. Each slide was parcellated into a set of 256×256 tiles and those that did not contain any tissue were automatically discarded. The remaining tiles were assigned with a ground-truth molecular profile (see **Online Methods: Biomarker acquisition**). **(b)**: The number of biomarkers per cancer type is shown as a heatmap. Biomarkers are grouped according to subclassification and cancer type.

### Pan-cancer predictability of multi-omic biomarkers from routine histology images with DL

We showed the overall feasibility of profiling multi-omic biomarkers, including clinically-relevant prognostic markers using histomorphological characteristics of standard H&E-stained WSIs. More than half of the models reached an AUC of 0.633 (**Figure 2a - left**). The observed AUC was greater than or equal to 0.711 for 25% of the models and above 0.829 for 5%. The top 1% models (n=135) returned an AUC of at least 0.904. For the majority of the biomarkers, the predictability was highly consistent with the standard deviation being less than 0.1 AUC, and the difference between the minimum and maximum performance being less than 0.2 AUC (**Figure 2a - right**). Most of the biomarkers showed better-than-random performance across all omics/biomarker types (**Figure 2b, Table 1**). The lowest average performance was seen in the prediction of metabolic pathways (AUC 0.564 ±0.081), and the highest performing models were from the standard clinical biomarkers (AUC 0.742 ±0.120). Variability across different models of the same biomarker type showed similar trends compared to that in the overall distribution, with the standard deviation for all omics being around 0.2 AUC and an increase in variance considering the range in minimum-maximum performance (**Figure 2c**).

**Figure 2.**
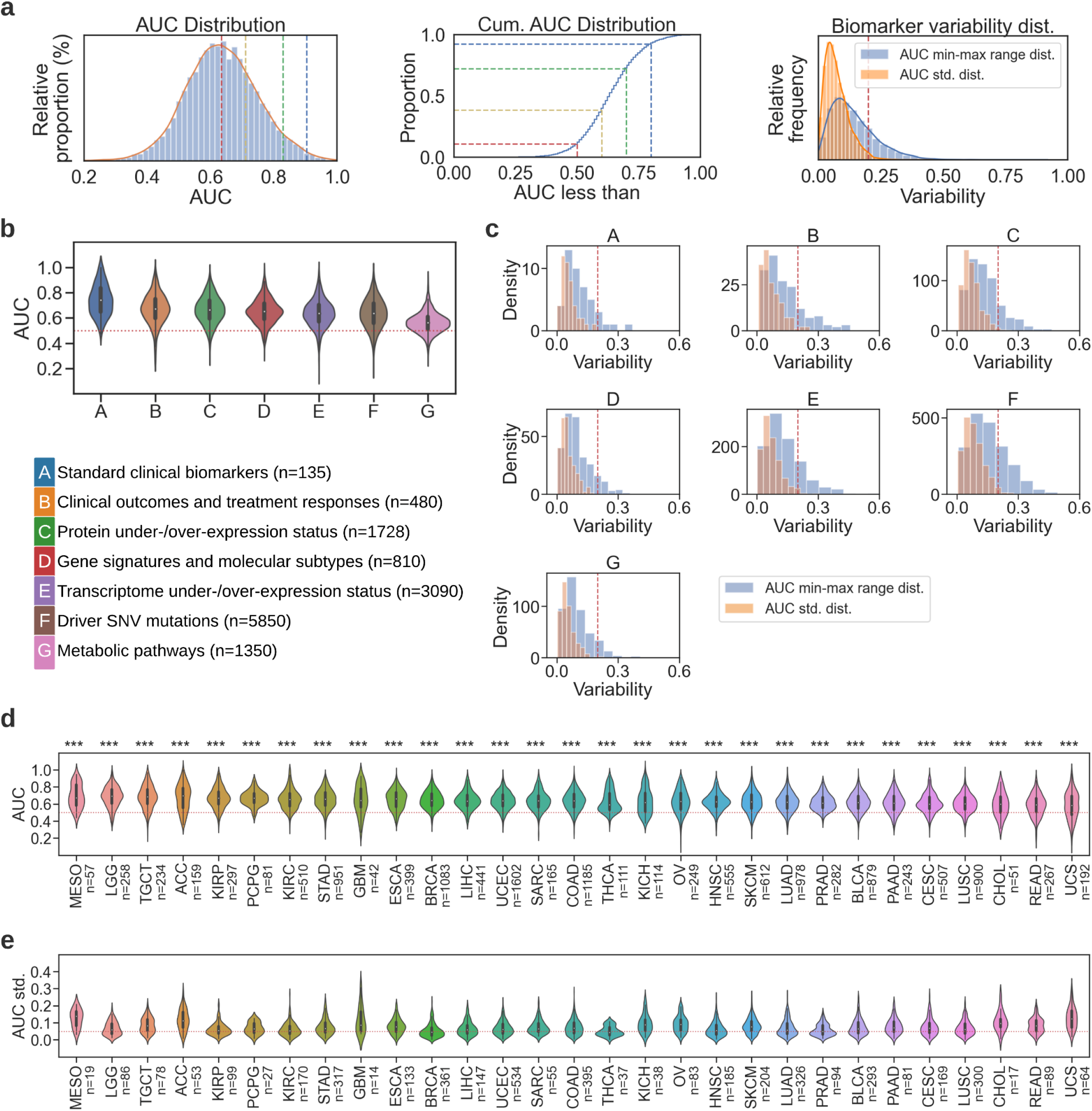
| Multi-omic biomarkers can be predicted directly from histo-morphology across multiple cancer types. (**a**): (*Left*) Histogram distribution and kernel density estimation of AUC values for all models (n=13,443). (*Middle*) Cumulative AUC distribution shows the proportion of models that have AUC less than the shown markers at 0.5, 0.6, 0.7, and 0.8. *(Right)*: Standard deviation and min-max range distribution of model performance in AUC. **(b)**: Violin plots showing the AUC distribution of each biomarker type. The same coding in **Table 1** was used to abbreviate the biomarker types. Box-plots were used to represent the data points in violins, with whiskers showing the 1.5x interquartile range and median values indicated with white dots. **(c)**: Standard deviation (orange histogram) and min-max range distribution (blue histogram) of model performance in AUC across the CV folds. **(d):** Violin plots showing the AUC distribution per cancer type (see **Extended Data Table 2** for cancer abbreviations). Plots are sorted by average intra-study AUC. DLBC, UVM, and THYM were excluded from this analysis due to only constituting one to seven valid targets across all biomarker types. Number of models per cancer type is given in prantheses. **(e):** Violin plots showing the standard deviation distribution of model performance across different folds of each biomarker (in the same order as in **(d)**. Number of standard deviation values per cancer type is given in parentheses.

**Table 1:** Average performance and standard deviation for all omic types, alongside their brief description. Number of biomarkers tested for each biomarker type is given in parentheses in the last column.

| Biomarker type / omic | Description | Mean AUC ± std. |
| --- | --- | --- |
| Standard clinical biomarkers | Established biomarkers that are routinely used in clinical management, as described in a related study<sup>11</sup>. | 0.742 ± 0.12<br>(n=135) |
| Clinical outcomes and treatment responses | Clinical outcomes such as overall survival and treatment responses to a therapy or drugs. | 0.671 ± 0.12<br>(n=480) |
| Under-/over-expression of proteins | Under- and/or over-expression status of proteins encoded by driver genes. | 0.666 ± 0.108<br>(n=1728) |
| Gene signatures and molecular subtypes | Biomarkers that are highly relevant for prognosis and targeted therapies, including molecular subtypes and gene expression signatures, as compiled in a related study<sup>11</sup>. | 0.653 ± 0.097<br>(n=810) |
| Under-/over-expression of driver genes | Under- and/or over-expression status in driver genes at the transcript level. | 0.637 ± 0.108<br>(n=3090) |
| Driver SNV mutations | SNV mutations in driver genes associated with FDA-approved therapies or known-relevancy for specific treatments. | 0.634 ± 0.117<br>(n=5850) |
| Metabolic pathways | Transcriptional activity in driver genes involved in metabolic pathways. | 0.564 ± 0.081<br>(n=1350) |

To show the performance of biomarker prediction for each cancer type, we plotted the distribution of AUC values for all the investigated malignancies with sufficient samples (**Figure 2d**) and provide the average performance with standard deviations in **Extended Data Table 1**. Overall, performance was significantly better than random across all cancer types (i.e. mean AUC > 0.5 and p < 1e-05 for all malignancies, where statistical significance for each cancer was measured with a two-sided t-test performed between the AUC values of all models belonging to a cancer type and a set of random AUC values with a mean of 0.5 and the same standard deviation as the compared AUC distribution). The lowest general performance was obtained in uterine carcinosarcoma (UCS) with a mean AUC of 0.586 (±0.158), and the highest performing models were in mesothelioma (MESO), where an average AUC of 0.693 (±0.137) was measured. Variability across CV folds of each biomarker was mostly stable, with standard deviations centring around 0.05 AUC in most of the studies (**Figure 2e**). Breakdown of the overall predictability performance for each malignancy depending on the type of biomarker is provided in **Extended Data Figure 1**.

### Feasibility of predicting genetic alterations from histology

Recent pan-cancer studies have shown that mutations can be detected from histomorphological features with DL^11, 14^. In our study, we extend the previous work on predicting mutations to a total of 1,950 genetic biomarkers. We focused on driver genes that are associated with disease-specific therapies approved by the Food and Drug Administration (FDA) or are known to be relevant for specific treatments based on evidence from clinical guidelines or well-powered studies with consensus from experts in the field^30^. In our experiments, we used the genomic profiles available from the TCGA project (**Online Methods: Biomarker acquisition**).

Genetic alterations were significantly predictable across most of the investigated cancer types (**Figure 3a**), with a mean AUC of 0.636 (±0.117). More than 40% of the mutations were detectable with an AUC of at least 0.65, and considering the highest performing mutations in each cancer type, almost all major malignancies had at least 10 mutations that were predictable with an AUC of 0.70 or above. Among them, endometrial carcinoma had the highest number of predictable mutations (n=112 out of 442 genes). It was followed by colon cancer (n=62 out of 250 genes), gastric cancer (n=58 out of 178 genes), skin melanoma (n=29 out of 109 genes), lung adenocarcinoma (n=28 out of 180 genes), and breast cancer (n=26 out of 128 genes). Among all the tested mutations, the top-performing ones were *NUMA1* and *JAK1* in clear cell renal cell carcinoma, *PDGFRB* and *BCL6* in lung cancer, *IRS2* in endometrial cancer, and *GNAS* in breast cancer, each with an AUC of at least 0.89. A large number of genes were highly predictable across multiple cancer types (**Extended Data Figure 3a**). Notably, alterations in *TP53* were detectable in many malignancies, with 7 out of 22 cancers tested having an AUC of at least 0.7 and 14 of them showing AUCs greater than 0.65, reaching up to 0.841 for brain lower grade glioma, up to 0.785 for breast cancer and up to 0.771 for endometrial cancer.

**Figure 3.**
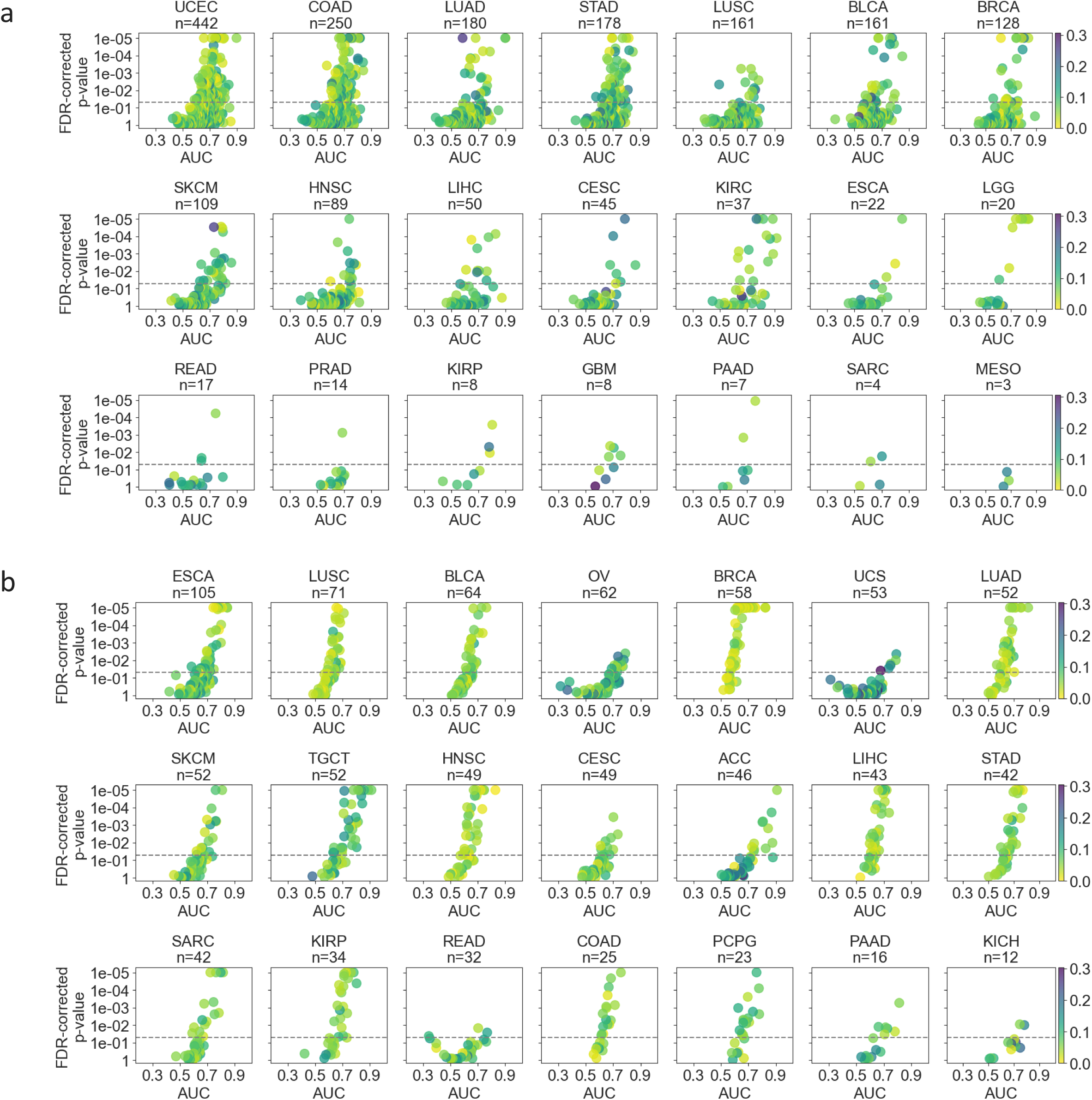
| DL could predict genetic alterations and transcriptome expression status from routine histology images across many cancer types. Scatter plots show the average test AUC for each model trained to predict **(a)** genetic alterations and **(b)** under-/over-expression status of transcriptomes across selected cancer types. Each marker in the scatter plot represents a tested biomarker. A two-sided t-test was applied to prediction scores of each model to assess the statistical significance and the corresponding p-values were corrected for false discovery rate (FDR). The y-axis of each plot was inverted and the p-values were log-transformed for visualisation purposes. p values smaller than 1e-05 were set to 1e-05 to avoid numerical errors during transformation. The statistical significance threshold of 0.05 is marked with a dashed line. Colour shading of the markers indicates the standard deviation of the predictability performance for each biomarker. Plots are ordered by the number of tested biomarkers in each row. Due to space limitations, cancer types with fewer biomarkers are not shown in this figure but are provided in **Extended Data Figure 2**. Cancer abbreviations are defined in **Extended Data Table 2**.

### Feasibility of inferring under-/over-expression of transcriptomes from diagnostic histology slides

Analysis of gene expressions is fundamental to better understanding the underlying cancer mechanisms and potentially help with improving cancer diagnosis and drug discovery^31^. While it is already known that genetic alterations could potentially be detected from histomorphology with DL, studies to understand the extent of predictability at transcript and protein levels have been rather limited. Recently, it has been shown that transcriptomic profiles are correlated with histomorphological features detected with DL in an annotation-free setup^15^. Our study took a more direct and comprehensive approach and trained deep learning models to predict the under- and/or over-expression status in selected driver genes, using the transcriptomic profiles available from TCGA. A total of 97 and 933 genes across multiple cancers were qualified to study the predictability of under- and over-expression at the transcript level, respectively (**Online Methods: Biomarker acquisition**).

The gene expression status was predictable across most of the investigated cancer types (**Figure 3b**) with a mean AUC of 0.637 (±0.108). The average performance was slightly lower for the under-expressed genes (mean AUC 0.633 ±0.115). Expression status in at least 40% of the genes was detectable with an AUC of 0.65 or above. Esophageal carcinoma and testis cancer were the malignancies with the highest number of genes with a predictable expression status (a total of 28 out of 105 and 52 genes, respectively), defined as an AUC level of at least 0.70. It was followed by ovarian cancer (18 out of 62 genes) and adrenocortical carcinoma (16 out of 46 genes). Almost all of the top-performing genes were that of over-expression, with *PMS2* in thymoma; *CARD11*, *LASP1*, *STIL, POLE, KMT2C*, and *CLIP1* in testis cancer; *ERC1, WRN, OLIG2, FANCC,* and *ACSL6* in adrenocortical carcinoma; and *SOX2* and *NDRG1* in esophageal carcinoma leading in performance with AUCs ranging 0.832-0.911. On the predictability of under-expressed genes, the most notable ones were *RHOA* in thymoma (AUC 0.908 ±0.05), *LSM14A*, *THRAP3*, and *MTOR* in the brain lower-grade glioma (AUCs ranging from 0.785 to 0.818) and *BAP1* in mesothelioma (AUC 0.818 ±0.084). Expression status of many genes were predictable across multiple cancer types (**Extended Data Figure 3b**).

### Feasibility of predicting protein expression level status with DL

As the next step in our multi-omics pan-cancer study, we assessed the ability of DL in detecting histomorphological changes that might be associated with the alterations in the expression of proteins. Towards this end, we trained models to predict the under- and/or over-expression status of proteins associated with certain driver genes based on proteomic profiles provided by TCGA. It is worth noting that association with a gene in this context refers to the encoding of a protein by that gene and we use “associated with/encoded by” interchangeably throughout the paper. A total of 267 and 309 driver genes were qualified to evaluate the predictability of their corresponding under- and over-expression status at protein level, respectively (**Online Methods: Biomarker acquisition**).

We achieved an average AUC of 0.666 (±0.107), with the under-expression status being slightly less predictable on average (mean AUC 0.662, ±0.105) compared to its over-expressed counterpart (mean AUC 0.669 ±0.109). The expression status prediction of almost all genes performed above random (**Figure 4a**), with more than half of them being detectable with an AUC of at least 0.65 and over 30% of them further achieving an AUC above 0.70. Breast cancer had the highest predictability rate, where the expression status of 37 out of 107 genes was detectable with an AUC of at least 0.7. It was followed by clear cell renal cell carcinoma and low-grade brain glioma (25 out of 51 and 76 genes, respectively). The expression level status of a large number of proteins encoded by driver genes was highly predictable, with *TFRC*, *ATM*, and *PIK3CA* in low-grade brain glioma; *NRAS*, *FOXO3*, *MYC*, and *TP53* in papillary renal cell carcinoma; *CDKN1B* in head and neck squamous cell carcinoma; and *MYC* in sarcoma, exhibiting the top performance with AUCs ranging 0.835-0.974. Multiple under-expressed proteins were also predictable to a great extent, the top-performing ones being *CASP8*, *MET*, *BCL2*, and *SETD2* in breast cancer; *AR* and *TFRC* in gastric cancer; and *VHL* in lung cancer with AUCs ranging 0.814-0.866. We found that the p53 protein over-expression (encoded by *TP53*) was consistently predictable in six of the eight tested cancers, including renal cell carcinomas, lower-grade brain glioma, and endometrial cancer, with AUCs ranging from 0.672 to 0.835. Expression status of many other proteins were also detectable across multiple cancer types (**Extended Data Figure 3c**).

**Figure 4.**
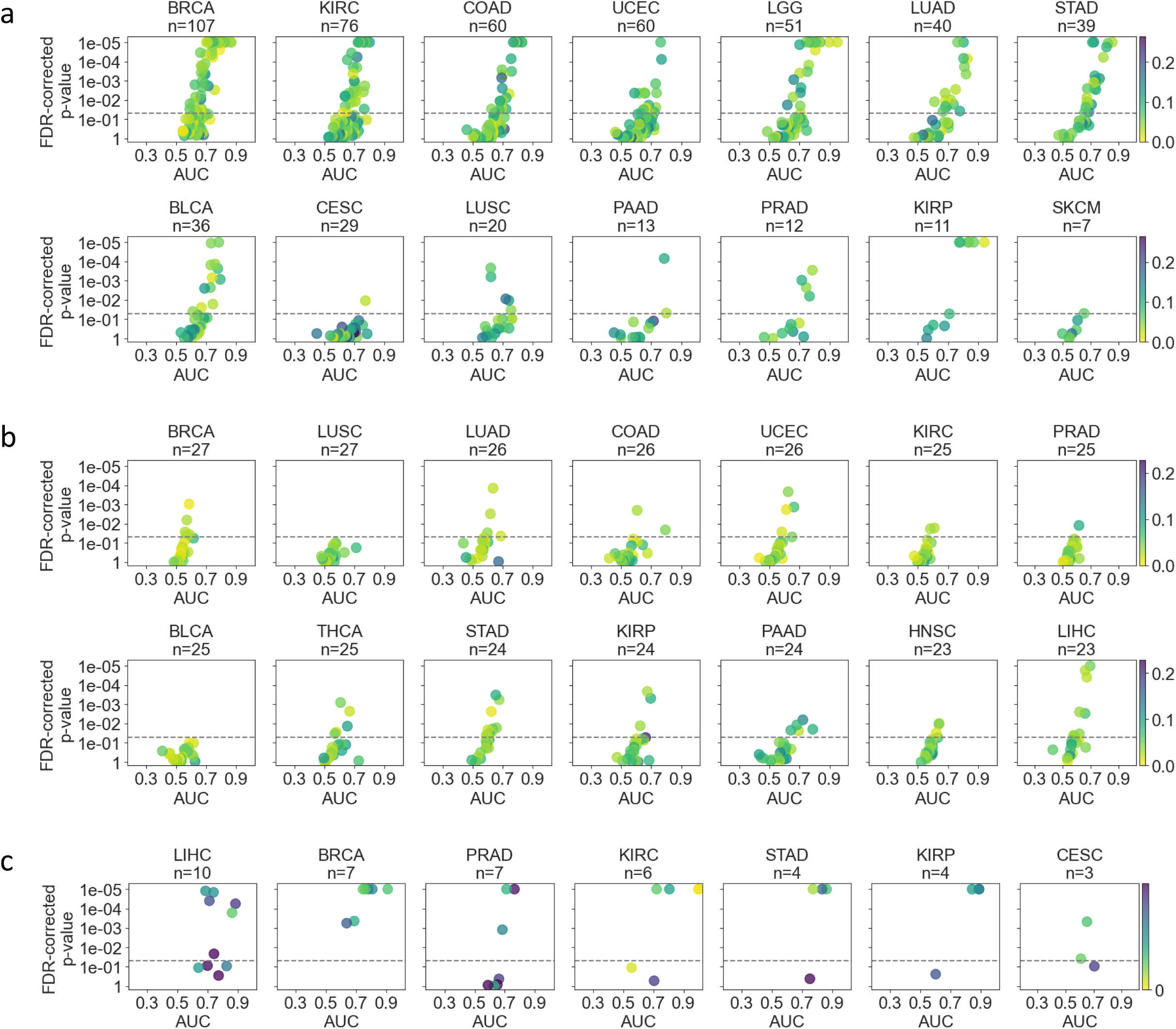
| Protein expression status, metabolic pathways and standard clinical biomarkers could be inferred with DL across many cancer types. Scatter plots show the average test AUC for each model trained to predict **(a)** under-/over-expression status of proteins, **(b)** metabolic pathways, and **(c)** standard-of-care biomarkers across selected cancer types. Plots are ordered by the number of tested biomarkers in each cancer type. Due to space limitations, cancer types with fewer biomarkers are not shown in this figure but are provided in Extended Data Figure 2. Please refer to the caption of Figure 3 for a detailed explanation of the visualisation.

### Predictability of transcriptomic and proteomic biomarkers is positively correlated

Despite the average predictability of the protein expression status being higher than that of the transcriptome (**Table 1**), both omic types had around 200 highly-predictable genes (i.e. AUC > 0.7) and are clustered around a comparable median. The slightly lower overall performance of transcriptome expression prediction may be attributed to it inherently being less predictable compared to its proteomic counterpart. Considering how predictability changes across the molecular landscape (**Extended Data Figure 4**), we measured a positive Pearson correlation of 0.227 (p<1e-05) and 0.124 (p=0.203) between the transcriptomic and proteomic biomarkers with regard to their under- and over-expression status, respectively. A positive linear relationship (Pearson correlation: 0.131, p<0.01) also existed between under-expressed transcriptomes and genetic alterations.

### Feasibility of inferring metabolic pathways from histology

We assessed the capability of DL for inferring transcriptional activity in driver genes associated with metabolic pathways (**Online Methods: Biomarker acquisition**) directly from histopathology as our final target in the omic landscape. Overall, metabolomic characteristics were less predictable than those of the other omic types, yielding an average AUC of 0.564 (± 0.081). Despite 87% of the pathways having a non-random performance (**Figure 4b**), only 9% of them were likely to be strongly predictable among a total of 450 tested targets (i.e. based on an AUC > 0.65). Considering the individual performance of each metabolic pathway, we found that nucleotide metabolism in testicular germ cell tumours and pancreatic cancer; nuclear transport in colon adenocarcinoma, skin cutaneous melanoma, and lung cancer; mitochondrial transport and histidine metabolism in ovarian cancer were highly predictable, with AUCs ranging from 0.720 to 0.847. Similarly, it was possible to predict certain pathways involved in beta-oxidation of fatty acids directly from histomorphology to a certain degree (AUC > 0.65) for another four cancers, including thyroid carcinoma, hepatocellular carcinoma, ovarian cancer, as well as adenocarcinomas of the pancreas and stomach.

### Feasibility of predicting standard clinical biomarkers with DL

We tested the feasibility of DL to predict the established biomarkers that are routinely used in clinical management. Towards this end, a set of standard of care features was compiled by following the biomarker acquisition approach in ^11^, including data for tumour grade, microsatellite instability (MSI) status (in colorectal and gastric cancer), histological subtypes, hormone receptor status (in breast cancer), and Gleason score (sum, in prostate cancer) (**Online Methods: Biomarker acquisition**).

Among all investigated omics and features, standard pathology biomarkers showed the highest predictability performance with an average AUC of 0.742 (±0.120). None of the biomarkers had a performance worse than random (i.e. all AUCs > 0.5) and almost 30% of them could be inferred with an AUC of above 0.8, a sign of very high predictability (**Figure 4c**). As expected, histological subtypes were in general highly predictable, especially for breast cancer, renal cell carcinomas, hepatocellular and gastric cancer. Prediction of molecular features in clear cell and chromophobe subtypes of renal cell carcinoma had the highest performance, reaching up to an AUC of 0.999. Invasive ductal carcinoma (IDC) and invasive lobular carcinoma (ILC) subtypes of breast cancer were well detectable from WSIs, with AUCs ranging 0.759-0.908. Our models were able to predict hormonal receptor status in breast cancer, with AUCs of 0.806 and 0.744, for oestrogen (ER) and progesterone (PR) receptors, respectively. Notably, multiple clinical biomarkers important for hepatocellular cancer could also be accurately inferred from histology, including growth patterns (AUC up to 0.862) and the etiological status of non-alcoholic fatty liver disease (NAFLD, AUC 0.826 ±0.054). Another highly predictable biomarker was MSI status, which was detectable in both colon and gastric cancer with average AUCs of 0.716 and 0.773.

### Feasibility of inferring molecular subtypes and gene expression signatures from routine images

To evaluate the capability of DL to detect molecular subtypes and gene expression signatures of cancer from WSIs, we compiled a set of well-established features with clinical and/or biological significance, by closely following the experimental details in a previous study^11^. This includes features that are relevant for prognosis and targeted therapies such as molecular subtypes and clusters, immune-related gene expressions, homologous recombination defects, cell proliferation, interferon-γ signalling and macrophage regulation and hypermethylation/mutation^32–34^ (**Online Methods: Biomarker acquisition**). Given their association with higher-level functions, these biomarkers may potentially have a larger impact on the morphology than the previously-assessed alterations, especially compared to single mutations^11^.

Overall, molecular subtypes and gene signatures were quite predictable with an average AUC of 0.653 (± 0.097). Almost half of them were detectable at an AUC level greater than 0.65 (**Figure 5a**). The most predictable biomarkers were observed in breast cancer (18 out of 28 biomarkers) and the adenocarcinomas of the stomach (16 out of 22 biomarkers) and colon (14 out of 24 biomarkers). Our method was capable of inferring TCGA molecular subtypes in multiple cancer types, including kidney renal papillary cell carcinoma (AUC up to 0.884 ±0.085), gastric cancer (AUC up to 0.875 ±0.048), lung squamous cell carcinoma (AUC up to 0.861 ± 0.015), and breast cancer (AUC up to 0.859 ± 0.028). Notably, the average predictability for PAM50 subtypes in breast cancer was 0.752 (± 0.080), reaching up to an AUC of 0.871 (±0.015) for the Basal subtype. Consensus molecular subtypes in colon cancer (i.e. CMS1, CMS2, CMS3, CMS4) were also highly detectable with an inter-subtype average AUC of 0.763 (±0.068), reaching up to 0.821 (±0.083) for CMS1. Cell proliferation and hyper-methylation were among well-predicted biomarkers, especially in cancers of breast, stomach, colon, and lung, with AUCs reaching up to 0.854.

**Figure 5.**
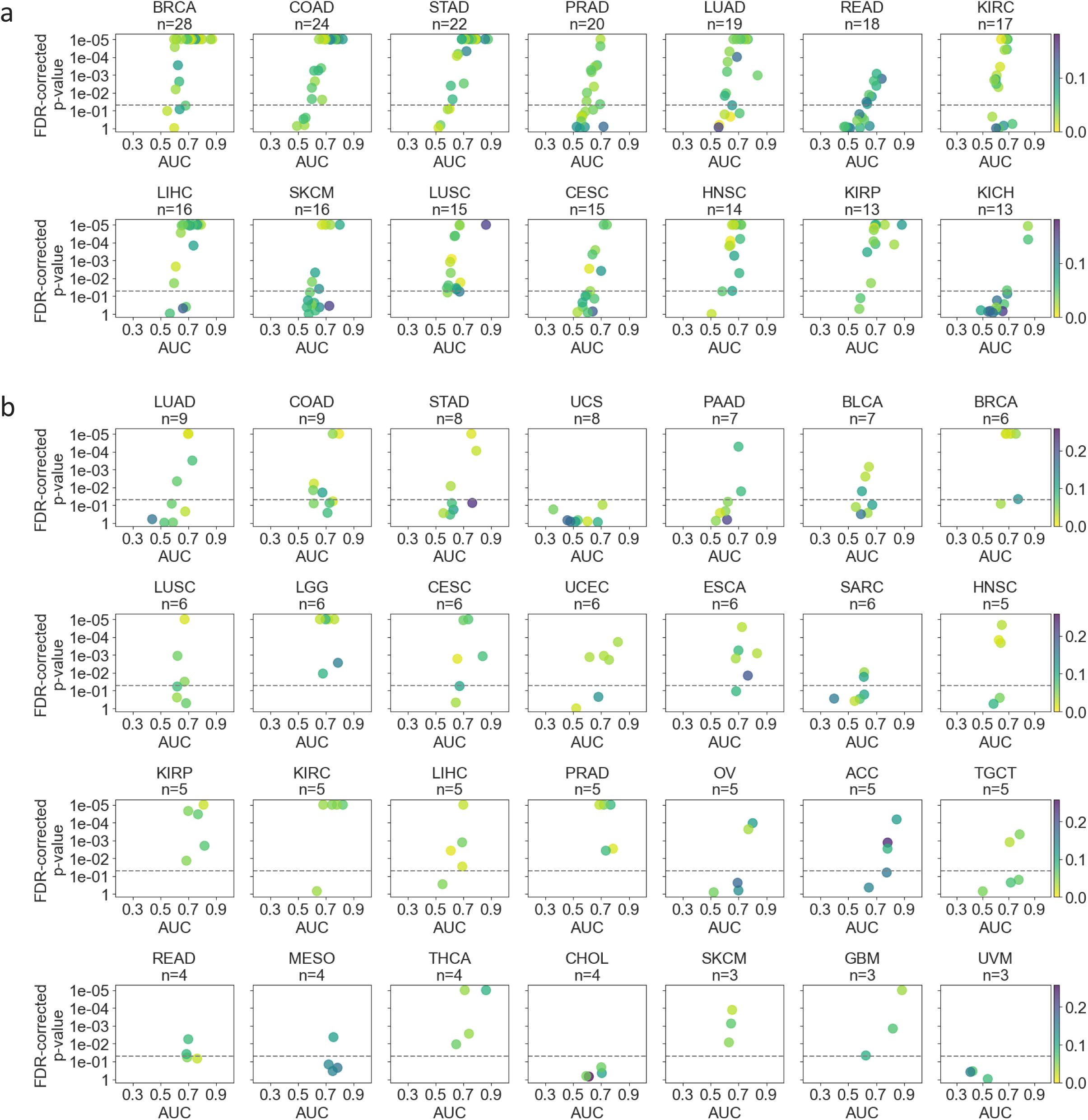
| DL could infer gene signatures, molecular subtypes, and clinical outcomes from diagnostic histology slides. Scatter plots show the average test AUC for each model trained to predict **(a)** gene signatures and molecular subtypes, and **(b)** clinical and treatment outcomes across selected cancer types. Plots are ordered by the number of tested biomarkers in each cancer type. Due to space limitations, cancer types with fewer biomarkers are not shown in this figure but are provided in **Extended Data Figure 2**. Please refer to the caption of **Figure 3** for a detailed explanation of the visualisation.

### Feasibility of inferring clinical outcomes and treatment response from diagnostic histology slides

Ability to accurately estimate prognosis can be vital for clinical management operations. Previous work has focused on developing prognostic models from routine clinical data, standard of care features, histopathological assessment, molecular profiling, and more recently, morphological features acquired via DL^35–39^. There have also been attempts to use ML approaches and image-based features to predict clinical endpoints in different cancers, such as melanoma and non–small cell lung cancer^40, 41^. In our study, we explored the end-to-end predictability of prognostic outcomes directly from histology across multiple cancer types by treating the clinical outcome endpoints such as overall survival (OS), disease-specific-survival (DSS), disease-free-interval (DFI), and progression-free interval (PFI) as potential prognostic biomarkers^42^. We further expanded our analysis towards detecting treatment responses directly from WSIs to assess whether DL models can correlate histomorphological features with the outcome of a therapy or drug (**Online Methods: Biomarker acquisition**). To the best of our knowledge, this study constitutes the first systematic attempt to assess the DL-based predictability of drug responses across multiple cancer types.

Overall, the predictive performance of clinical outcomes and treatment responses was relatively high compared to other biomarker types and omics, with a mean AUC of 0.671 (±0.12). Almost 40% of the tested targets were predictable at an AUC level of 0.70 or above (**Figure 5b**). We acquired the best overall performance in glioblastoma multiforme, adrenocortical carcinoma, and kidney chromophobe with mean AUCs of 0.77. They were followed by renal papillary cell carcinoma, mesothelioma, thyroid carcinoma, prostate cancer, renal clear cell carcinoma, and esophageal carcinoma, with overall AUCs ranging from 0.731 to 0.76. *OS* in kidney chromophobe, glioblastoma multiforme, thyroid carcinoma, and adrenocortical carcinoma; DFI in esophageal carcinoma and renal clear cell carcinoma; DSS in glioblastoma multiforme; residual tumour status in endometrial carcinoma, and the treatment response in papillary renal cell carcinoma were among the top-performing targets with AUCs ranging from 0.815 to 0.924.

Among the 20 drugs we investigated in our study, DL was able to predict the response in half of them with an AUC of at least 0.7. Cisplatin was the most notable drug with AUCs ranging 0.763-0.837 in cervical, testis and gastric cancers. The other predictable drugs were temozolomide in lower-grade glioma; paclitaxel in breast cancer; leucovorin, oxaliplatin, and fluorouracil in colon cancer; etoposide in testis cancer; and gemcitabine in pancreatic cancer.

### Pan-cancer predictability is consistent across different datasets

To show that a comparable performance can be achieved across different datasets, we repeated our experiments using publicly available data from the Clinical Proteomic Tumour Analysis Consortium (CPTAC) (**Online Methods: External dataset**). We limited our analysis to the prediction of driver SNV mutations due to both TCGA and CPTAC relying on the same set of driver genes, hence exhibiting a relatively large overlap (**Online Methods: Biomarker acquisition).** A total of 176 driver genes across seven cancer types were identified that have available mutation data in both datasets. The investigated cancers were endometrial carcinoma (n=61 out of 176 equivalent biomarkers), pancreatic ductal adenocarcinoma (n=4), lung squamous cell carcinoma (n=21), lung adenocarcinoma (n=14), head and neck cancer (n=12), glioblastoma multiforme (n=4), and colon adenocarcinoma (n=60). A comparable overall performance was observed on both datasets across almost all tested cancer types (**Extended Data Figure 5**, p > 0.05 under two-sided t-test for all but COAD) with within-cancer average AUCs ranging 0.578-0.655 and 0.567-0.672 for TCGA and CPTAC, respectively. Performance across biomarkers varied more for CPTAC cohorts, as indicated by the shape of violin plots. Overall, our results indicate that the predictability of the biomarkers is consistent and dataset-independent.

## Discussion

This study assessed the general feasibility of predicting a plethora of biomarkers from a pan-cancer perspective, using DL and histomorphological features extracted from H&E-stained diagnostic slides. Morphological visual characteristics captured from histology with an encoding-decoding DL model were used for inferring biomarkers across a wide omics spectrum with varying performance depending on the biomarker type. We observed relatively high predictability for certain genetic alterations (e.g. *TP53*) and especially for clinically-relevant markers (e.g. standard-of-care features and molecular subtypes) across multiple cancer types. The prediction of a biomarker appeared to be consistent, with the cross-validated models performing similarly across different subsets of the sample space. A comparable predictive performance was obtained for certain biomarkers when experiments were repeated on an independent dataset, further showing the overall capability of DL for molecular profiling across multiple cancer types.

The performance of a biomarker did not seem to depend on the factors intrinsic to the tested populations, such as the number of cases (i.e. population size) and the proportion of positive-negative samples in the whole population (**Extended Data Figure 6a**). We argue that the dataset size is more important for the stability of the results, rather than the performance itself, i.e. having a larger training subset is likely to yield models with more consistent performance. In addition, our biomarkers typically had an unbalanced distribution, due to most of the biomarkers having a much smaller number of positive cases than that of negative. We tackled this problem at training time by oversampling more from the underrepresented class and later assessed its impact on the stability of performance. Towards this end, we compared the sample size and class ratios to the range of AUCs across cross-validation folds for each biomarker (as measured by standard deviation), and observed a negative relationship for both factors (**Extended Data Figure 6b**). This indicates that the performance becomes less variable with an increasing number of samples and a more balanced dataset. We further assessed the importance of other factors such as the type of cancer and biomarker class for predictability (**Extended Data Figure 7**), and observed that the combined impact of biomarker types was larger than that of cancer types. In addition, we performed a correlation analysis to measure the relationship between the biomarker predictability and tumour purity (**Extended Data Figure 10**). We found that there was no strong relationship between the two variables. This might indicate that tumour purity is unlikely to impact the predictability.

**Figure 6.**
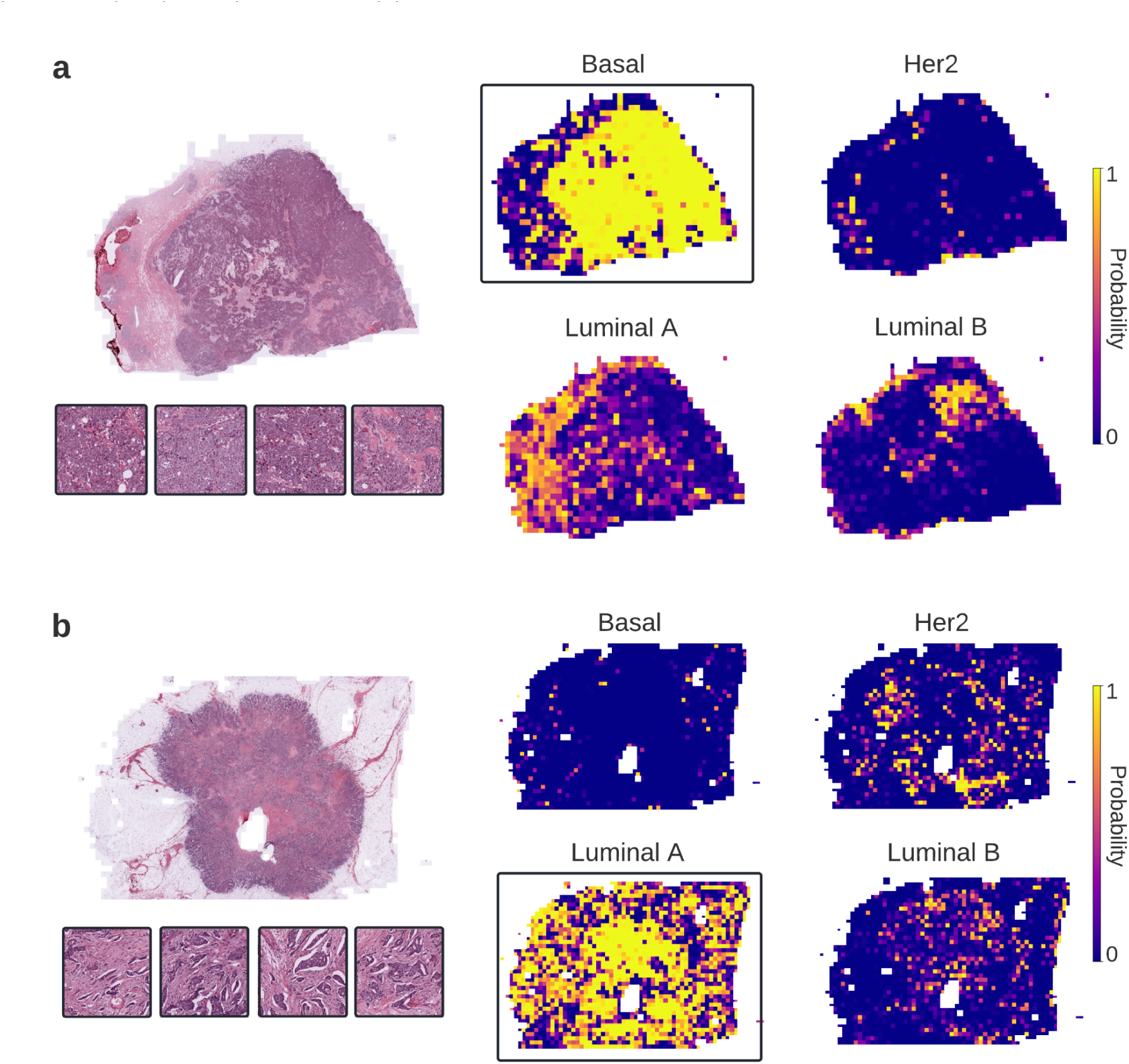
| Visualisation of predictability with DL from histopathological images. **(a-b)** DL-based predictions for the molecular subtypes of breast cancer (i.e. Basal, Her2, Luminal A, and Luminal B) are visualised for two selected patients using heatmaps. The correctly-predicted subtype in each case is enclosed with a rectangle and the highest ranking tiles from that class are given alongside the original WSI. The Basal type **(a)** shows sheets of tumour cells without any discernible gland formation, while the Luminal A patient’s tumour **(b)** is composed of well formed glands. Considering the heatmaps in both cases, one can notice that DL models can identify spatial regions that are relevant for the target class.

In our study, we found that detecting alterations from histology at different levels of the omics landscape was mostly feasible for the majority of the investigated genes. Certain mutations likely to be associated with poor clinical outcomes were highly predictable across multiple cancer types, notable examples being those harboured by *TP53, BAP1, MTOR,* and *GNAS*. For *TP53*, the relatively high and consistent predictability might be attributed to the tumours with *TP53* mutations likely being poorly-differentiated and can be linked to higher grade cell changes, which are visually discernible^14^. Identifying patients with certain alterations is highly important for precision treatment and can pave the way to develop targeted therapies. For instance, recent studies show that detection of mutations in *BAP1* can potentially be useful for the development of targeted treatment strategies in clear cell renal cell carcinoma^43^. Similarly, *MTOR* mutations can serve as biomarkers for predicting tumour responses to mTOR inhibitors, which are already being used to treat human cancer^44^. One of the highly predicted genes, *GNAS,* is known to promote cell proliferation and migration in breast cancer when expressed at high levels, and thus, can potentially be used as a therapeutic target^45^.

Identification of downstream changes in tumours and how they differentiate from normal cells can help uncover the complex mechanisms that control the anticancer drug response and potentially yield better prediction of therapeutic outcomes^46–49^. Despite many studies targeting tumour metabolism, assessing the detectability of transcriptomic, proteomic, and metabolic activities from WSIs has been highly limited^15, 50, 51^. In our study, it was possible to predict the under-/over-expression status of transcriptomes and proteins to a certain degree. For instance, we found that p53 over-expression was consistently predictable across multiple cancer types. Compared to the other omics of the molecular landscape, a relatively lower overall predictability was found for metabolic pathways. A previous study also reported that the predictability of pathway alterations could be lower than that of altered genes, suggesting that the molecular changes at gene level are likely to be responsible for the morphological patterns that drive the predictability^29^. In our study, certain metabolic pathways that are dysregulated in tumours, such as the nuclear transport pathway and fatty acid beta-oxidation, were detectable from histology in several cancer types^52, 53^. Another highly predictable pathway was nucleotide metabolism, which is known to support the uncontrolled growth of tumours when increased^54^. This presents a promising opportunity in potentially utilising metabolite markers in routine clinical uses.

Our findings indicated that DL can infer well-established standard-of-care clinical biomarkers, gene expression signatures and molecular subtypes from histopathological images. Our results mostly agree with those obtained in the recent pan-cancer study by Kather et al.^11^. Given the similarities between the two studies, our results can confirm their findings and provide more comprehensive evidence for the feasibility of detecting molecular biomarkers from histology.

Classification of residual tumours is a critical stage for the course of treatment and is considered an important prognostic biomarker^55^. In our analysis, we found that DL could detect the occurrence (or lack thereof) of residual tumours in multiple cancers, which might indicate that some visual clues correlated with complete remission after treatment are already present in histomorphology at the time of diagnosis. While other clinical outcomes, such as overall survival (OS) and disease-specific survival (DSS), were also predictable to a certain extent, one should note that the definition of clinical outcomes may not always be accurate, especially for the cancer types that need longer follow-up times, have a small cohort size or have a limited number of events^42^. This study assessed the predictability of drug responses from H&E-stained images and revealed that the complete response to a number of drugs such as cisplatin, temozolomide and paclitaxel could be detected with DL. These findings might indicate the potential of DL approaches for precision medicine, allowing oncologists to select a treatment that could work best for the patients by only analysing routine histology slides.

One of the limitations in our study is that we imposed certain constraints during biomarker acquisition to reliably assess the predictability. For instance, multi-omic biomarker profiles were only computed for driver genes. Similarly for genetic alterations, only single variant mutations were considered. We also made certain assumptions in case the actual molecular profiles of a biomarker type were not available. For example, metabolic biomarkers were indirectly derived from the transcriptional activity in driver genes involved in metabolic pathways, so they can be considered as a proxy biomarker rather than reflecting the real pathways. This may explain the reason for suboptimal results obtained with metabolomic biomarkers. On the other hand, a recent study has shown that prediction performance at the single gene level is likely to be better than that for pathway alterations^29^. This may indicate that individual genes leave a stronger morphological signature than genetic pathways in H&E-stained images. Given this, we may also expect transcriptomic activity to be more detectable than of composite pathways, but this raises the need for further research. Another limitation of the study comes directly from the data itself. While we compared predictability across multiple cancers and omic types, all biomarkers were tested under different sample size and prevalence conditions. Many potential biomarkers were simply discarded during data acquisition, as they did not contain enough positive samples. It is also worth noting that TCGA is known to have site-specific fingerprints inherent in digital slides which may bias the accuracy of predictive models and one may consider using more sophisticated splitting techniques to avoid mixing samples from the same sites into training and validation sets^56^. We assessed the predictability based on AUC, the most commonly used evaluation metric for predictive models under the presence of class imbalance. However, in scenarios where only a few examples are available for the minority class (such as rare mutations), AUC values can be less reliable. Performance as estimated by AUC can drastically change based on correct and incorrect predictions of the minority class. One should note this inherent drawback when interpreting the results of biomarkers with very small sample size and low prevalence.

Exploring the “black-box” representation of DL models can be useful to reveal the morphological patterns that may be linked to certain alterations or phenotypic outcomes^11, 14^. One way to visualise the spatial regions that are critical for inferring a biomarker status is superimposing the tile-level prediction scores (i.e. probabilities) onto WSIs to create spatial heatmaps (**Figure 6**). The highest-ranking tiles within these heatmaps represent the visual characteristics learnt by the DL model to solve the prediction task at hand. For instance, different types of breast tumours may show distinct differences in morphology, which can be identified by DL and utilised to differentiate specific subtypes (**Figure 6a-b**). Similarly, top-ranked tiles from consensus molecular subtypes (CMS) of colon cancer (**Extended Data Figure 9a**) show distinct morphological patterns that consistently align with the histopathology of CMS subclasses as shown in previous studies^11, 19^. In addition, morphological traits associated with MSI unstable cases, e.g. containing large amounts of tumour-infiltrating lymphocytes, can be seen in the highest-ranking MSI tiles acquired from patients with colon and gastric cancers (**Extended Data Figure 9b-c**). While DL can identify clinically-relevant morphological features, it may also be useful to trace back the visual patterns that are associated with molecular alterations. For instance, highly-predicted tiles from a mutated *BRAF* case in papillary thyroid carcinoma show distinct histological features compared to its wild-type counterpart (**Extended Data Figure 9d**). In addition, DL may be a useful tool to explore visual characteristics with unknown links to histomorphology. For instance, no distinct features are known to distinguish the mutated *TP53* and its wild-type in breast cancer, but DL can still pick tumour tiles that show different visual characteristics for both classes (**Extended Data Figure 9e**). This may provide insights to better understand the potential impact of this alteration on cancer cell morphology.

Showing the overall feasibility of predicting multi-omic biomarkers with DL marks an important step in pursuit of achieving end-to-end detection systems from histology, which can potentially assist clinicians in patient management, accelerate diagnosis, and help develop more patient-centric treatments. The approach also presents an opportunity to apply multi-omic biomarkers in a rapid and low-resource setting without requiring time consuming and expensive biological tests. While this study has elucidated early observations on the factors determining a biomarker’s predictability, further understanding would be necessary before the mainstream adoption of DL-based methods for multi-omic biomarker profiling from standard tissue imaging in day-to-day clinical settings^57^. Our study showed that genetic alterations are detectable at different levels of the omics landscape for many genes. However, the benefit of integrating multi-omic data for predicting mutations is still not fully understood and further analysis is required to explore the impact of combining multiple omics on predictability^58^. The questions around the specific mechanism of biomarker detectability, such as the predictive pattern and how it is conserved across the different landscapes of the molecular landscape, merit more investigation. Future work will explore the internal representations of the predictive models to identify any associations between omics, tumour morphology and the model predictions. This study formulates the problem of predicting biomarker status from H&E-stained images as a single-task classification problem, however future studies should investigate explicit multi-task methods considering that certain biomarkers may be correlated with each other. This may potentially improve predictability performance.

## Online Methods

### Dataset

We conducted our experiments on the data provided from The Cancer Genome Atlas (TCGA) project, which was retrieved via the Genomic Data Commons (GDC) Portal (https://portal.gdc.cancer.gov/). The TCGA dataset consisted of 10,954 hematoxylin and eosin (H&E)-stained, formalin-fixed, and paraffin-embedded (FFPE) whole slides images of 8,890 patients, acquired from the following studies: breast cancer (BRCA), cervical squamous cell carcinoma (CESC), kidney renal papillary cell carcinoma (KIRP), clear cell renal cell carcinoma (KIRC), kidney chromophobe (KICH), skin cutaneous melanoma (SKCM), sarcoma (SARC), pancreatic adenocarcinoma (PAAD), ovarian serous cystadenocarcinoma (OV), prostate adenocarcinoma (PRAD), bladder urothelial carcinoma (BLCA), esophageal carcinoma (ESCA), thyroid carcinoma (THCA), lymphoid neoplasm diffuse large B-cell Lymphoma (DLBC), brain lower grade glioma (LGG), thymoma (THYM), head and neck squamous cell carcinoma (HNSC), uterine corpus endometrial carcinoma (UCEC), glioblastoma multiforme (GBM), cholangiocarcinoma (CHOL), liver hepatocellular carcinoma (LIHC), stomach adenocarcinoma (STAD), lung adenocarcinoma (LUAD), and lung squamous cell carcinoma (LUSC), colon adenocarcinoma (COAD), rectum adenocarcinoma (READ), adrenocortical carcinoma (ACC), mesothelioma (MESO), pheochromocytoma and paraganglioma (PCPG), testicular germ cell tumours (TGCT), uterine carcinosarcoma (UCS) and uveal melanoma (UVM). Only images scanned at a resolution of 0.5 microns per pixel (MPP) were kept and images with no MPP information were discarded, ensuring consistent resolution within each cancer cohort. The number of the images and patients included in the TCGA cohort are provided in **Extended Data Table 2**. DLBC, UVM, and THYM were excluded from certain results due to having less than seven valid targets considering all biomarker types. See **Biomarker acquisition** for more details on the biomarker inclusion criteria for each omic/biomarker type).

### External dataset

In order to assess the general feasibility of biomarker predictability on an external dataset, we repeated our experiments with the Clinical Proteomic Tumour Analysis Consortium (CPTAC) data, retrieved via the Cancer Imaging Archive (https://wiki.cancerimagingarchive.net/display/Public/CPTAC+Imaging+Proteomics). The CPTAC dataset consisted of 3,481 H&E stained images corresponding to 1,329 patients, acquired from the following seven different studies: lung adenocarcinoma (LUAD), colon adenocarcinoma (COAD), head-and-neck cancer (HNSCC), lung squamous cell carcinoma (LSCC), pancreatic ductal adenocarcinoma (PDA), glioblastoma multiforme (GBM), and uterine corpus endometrial carcinoma (UCEC). LUAD, GBM, and COAD were obtained from frozen tissues, whereas the rest of the datasets contained FFPE slides. Images with a resolution different than 0.25 MPP in COAD and 0.5 MPP in other datasets were discarded, ensuring consistent resolution within each cancer cohort. The details of the final images and patients included in the CPTAC cohort are provided in **Extended Data Table 3**.

### Ethics oversight

Only retrospective data was used in this study. The authors had no role in the recruitment of participants. Ethics oversight of the TCGA study is described at https://www.cancer.gov/about-nci/organization/ccg/research/structural-genomics/tcga/history/polic <u>ies</u>.

### Biomarker acquisition

#### Acquisition of actionable driver genes

Clinically relevant driver genes were retrieved from https://cancervariants.org59. We only considered driver genes that are known to associate with 1) FDA-approved disease-specific therapies and 2) response or resistance to therapies as shown in professional guidelines and/or based on well-powered studies with consensus from experts in the field based on evidence provided in another study^30^. Driver mutation and drug-associated data were acquired from the following sources: BRCA exchange, the Cancer Genome Interpreter Cancer Biomarkers Database (CGI), Clinical Interpretation of Variants in Cancer (CIViC), Jackson Laboratory Clinical Knowledgebase (JAX-CKB), the Precision Medicine Knowledgebase (PMKB). These source files already contained the associations between SNP mutations and phenotypes, allowing an expert pathologist to map them to TCGA studies. Finally using this mapping and driver mutation data per phenotype, we created a set of driver genes per TCGA study and subsequently used them to filter actionable biomarkers for transcriptomic, proteomic and genomic data.

#### TCGA genomic biomarker profiles

Genomic biomarker data was collected using the cBioPortal web API (https://docs.cbioportal.org/6.-web-api-and-clients/api-and-api-clients) and the GDC API (https://gdc.cancer.gov/developers/gdc-application-programming-interface-api). For each TCGA study, we retrieved all samples with associated diagnostic slides. The samples that did not have whole-genome or whole-exome sequencing data were excluded from further consideration, allowing us to assume that all genes of interest were profiled across the remaining ones. While there existed samples without WGS or WXS data with mutations, it was not possible to assume that genes with no mutations were present in their wild-type, as they might simply have not been sequenced. For all TCGA studies listed on the cBio portal, we acquired molecular profiles with the *MUTATION_EXTENDED* alteration type and all mutations belonging to these molecular profiles within the collected samples were retrieved and stored in an intermediate format. Finally, we created molecular profiles for all driver genes using this mutation data. A sample was considered positive for a driver gene if it contained at least one single nucleotide variant (SNV). SNVs are typically insertions, replacements or deletions of one base, but on a few occasions can be of multiple bases (e.g. “T” being replaced by “CGC”). The resulting profiles were filtered to exclude driver genes that had less than 10 positive samples in a given cancer.

#### Transcriptomic and proteomic profiles

Transcriptomic and proteomic data for TCGA datasets were retrieved from the cBioPortal API (http://www.cbioportal.org/api). cBioPortal provides z-scores, which were originally computed from the raw FPKM counts of gene expression and the corresponding number of standard deviations to the mean of expression values. These z-scores were acquired for each coding gene and each sample with an associated tissue slide in the TCGA studies. cBioPortal restricted the transcriptome z-score calculations to the samples in which the tumour comprised diploid cells. Proteomic z-scores were calculated among all available samples for a given cancer. The z-scores were binarised for each gene and sample based on thresholds chosen as follows: For each sample, genes with a z-score of less than or equal to *t_under* were considered *underexpressed* and those with a z-score of larger than or equal to *t_over* were considered *overexpressed*. We set {*t_under*, *t_over*} to {-2, 2} and {-1.5, 1.5}, for transcriptomic and proteomic data, respectively, based on their ability to divide the total distribution of z-scores into balanced classes over all genes of interest. These thresholds were then used to generate two under-/over-expression profiles for proteomic and transcriptomic genes. In an “under-expression” profile all samples that were considered underexpressed were labelled as positive whereas all other samples were labelled as negative. Similarly, overexpressed samples in an “over-expression” profile were assigned a positive label, while the remaining samples were considered negative. Finally, to reduce the number of target biomarkers, we limited the over-and under-expression profiles to only include the driver genes (see **Acquisition of actionable driver genes**) for each study. Furthermore, profiles that did not contain enough positive samples were excluded. The minimum number of positive samples for proteomic genes was set to 20. For the transcriptomic profiles, only the ones with at least a positive ratio of 10% and having a minimum of 10 positive samples were kept.

#### Metabolic pathways

Metabolic data was downloaded from https://www.ebi.ac.uk/biomodels/pdgsmm/index. It contained personalised genome-scale metabolic models (GSMMs) stored in XML format for 21 of the TCGA studies. We used the pathology atlas of the human cancer transcriptome data^60^ to acquire the generalised base network used to create the GSMMs and followed the same enrichment analysis as the original study^60^. The analysis included the following steps: The genes associated with a specific metabolic pathway were defined according to the gene-reaction relationship from the generic GSMM for human cancer. A gene set was considered enriched in a specific metabolic pathway if it significantly overlapped with the genes associated with the metabolic pathway using a hypergeometric test. Similarly, we derived a biomarker by comparing the genes present in the pathway for a sample, to all genes present in the generalised underlying network for the given pathway. We created sets of genes present for each pair of pathways/samples and performed t-tests comparing them to the set of genes for the same pathways in the generalised network. The resulting p-values were adjusted for multiple testing using the Benjamini-Hochberg procedure^61^ with an FDR rate of 0.05. All samples with an adjusted p-value of less than 0.05 were considered significantly associated with the pathway. We limited ourselves to a set of “pathways of interest” which contained 32 pathways selected based on the entropy of their positive/negative distributions over all studies, as well as a few hand-picked pathways with known connections to cancer phenotypes. We created biomarker profiles across 21 TCGA projects, by considering each pathway as a biomarker, where a sample was labelled as positive if the genes present for a pathway in the sample’s GSMM were significantly overlapping with the ones present in the generalised base network. Pathways constituting less than 20 positive samples in each study were excluded from the resulting profiles.

#### Standard of care features, gene expression signatures and molecular subtypes

A publicly available dataset provided as part of a relevant study on detecting clinically actionable molecular alterations was used to acquire the biomarker profiles for gene expression signatures, molecular subtypes and standard clinical biomarkers^11^. The dataset was originally curated from the results of systematic studies using the TCGA data (https://portal.gdc.cancer.gov/)32–34 and contained profiles for 17 TCGA datasets (please refer to the original study^11^ for the description of biomarkers and other details regarding the acquisition protocol). For certain biomarkers, we used the consensus opinion to map the molecular status to binary labels. For instance, considering microsatellite instability (MSI), all patients defined as MSI-H were included in the positive class, while microsatellite stable (MSS) and MSI-L patients were labelled as negative. Profiles with multiple categorical values were binarised with one-hot-encoding, where a profile was created for each category with only the samples of that category being set to positive. Non-categorical profiles with continuous values were binarised at mean after eliminating NaN values^11^.

#### Clinical outcomes and treatment responses

Survival data was acquired from the TCGA Pan-Cancer Clinical Data Resource (TCGA-CDR), a publicly available dataset that provides four major clinical outcome endpoints^42^, namely, overall survival (OS), disease-specific-survival (DSS), disease-free-interval (DFI), and progression-free interval (PFI). These endpoints were systematically binarized into actionable events by considering multiple clinical and prognostic features acquired from TCGA’s routinely-collected clinical data (https://portal.gdc.cancer.gov/) such as vital status, tumour status, cause of death, new tumour events, local recurrence and distract metastasis. The details of the integration of the clinical data into actionable survival outcomes are given in the original study^42^. Additionally, we added the residual tumour status acquired from the TCGA clinical files as another prognostic target. Patients with microscopic or macroscopic residual tumours (R1 or R2) were classified as positive whereas those with no residual tumour (R0) were included in the negative class^55^. Since TCGA-BRCA did not have residual tumour information, we used “margin_status”. “treatment_outcome_first_course” was used to create binary targets representing the treatment response. Towards this end, any patient with “Complete Remission/Response” was included in the positive class whereas “Stable Disease”, “Partial Response” and “Progressive Disease” were considered negative. Finally, clinical drug files in the TCGA datasets were used to identify drug responses. This was achieved by first unifying drug names based on the data provided in another study^62^ and then identifying drug-study pairs with enough samples. Finally, the “treatment_best_response” attribute was used to map the drug responses to binary categories, with “Complete Response” constituting the positive class and the others being negative. For both treatment and drug responses, we only focus on assessing the predictability of complete response from histology and classify other outcomes including partial response in the negative class.

#### CPTAC genomic biomarker profiles

Molecular profiling data was collected using the GDC API (https://gdc.cancer.gov/developers/gdc-application-programming-interface-api). We focused on the “Single Nucleotide Variation” (SNV) files, which contained information about mutations associated with substitutions of a single DNA base or deletion and insertion of a small amount of bases. The SNV files were retrieved using the files endpoint of the GDC API, with the following filters: *files.data_category = [“Simple Nucleotide Variation”]*, *files.data_type = [“Masked Somatic Mutation”]* and *files.experimental_strategy = [“WXS”]*. After acquiring the SNV files, it was possible to obtain a list of SNV gene mutations for each sample of the CPTAC dataset. Since the mutation data was based on whole exome sequencing, a gene was considered wild-type if it was not listed in the SNV file of a given sample. Finally, biomarker profiles were limited to only include the driver genes (see **Acquisition of actionable driver genes**). A sample was considered positive, if the sample contained at least one mutation for a given driver gene. The resulting profiles were filtered to exclude genes that had less than 10 positive samples in a given dataset.

### Experimental setup

We assessed the predictive performance of each biomarker in a 3-fold cross-validation setting, where the cases with a valid biomarker status in each dataset were split into three random partitions (folds), each having approximately the same proportion of positive samples. We trained and tested three models per biomarker, each time keeping aside a different fold for validation and using the remaining ones for training. This setting ensured that the predictability can be assessed on a different (held-out) validation set for multiple times, and consequently allowing us to assess the variability of model performance. The images were partitioned at the patient level so that no patient could appear in multiple folds. A biomarker profile with less than 10 positive patients was discarded from the study.

### Pre-processing pipeline and training details

A convolutional neural network (CNN) was used for predicting molecular profiles from H&E images as illustrated in **Figure 1**. A single CNN was end-to-end trained from scratch for each biomarker and fold, yielding a total of 13,443 unique models that were used to obtain the results presented in this study. Each model was trained on a set of 256×256 tiles acquired from H&E-stained WSIs, considering the whole histological material. A standard deviation filter was used to eliminate the tiles which do not contain any relevant information, allowing us to extract the tissue from the rest of the image. A slide was discarded from analysis if it contained fewer than 10 tiles after the filtering process. Macenko colour and brightness normalisation^63^ was applied to the remaining tiles before they were assigned with a ground-truth molecular profile (see **Biomarker acquisition**).

A CNN consisted of a feature extractor (encoder), a decoder and a classification module. The encoder can capture the tissue properties within tiles throughout a set of convolutional filters applied to tiles at various layers of depth, effectively encoding the high-level visual features into a d-dimensional feature vector, where *d* depends on the architecture of the CNN. These vectors are regarded as the fingerprints of the tiles and are submitted to both the decoder and the classification module. The decoder module takes a *d*-dimensional embedding as input and returns an output of the same shape as the original tile that the embedding represents. It consists of a series of transposed convolutional and upsampling layers that is used to reconstruct the original tile from the latent vector to achieve better representations of each tile that do not contain irrelevant features. The output of the decoder is compared against the original tile with mean squared error (MSE), or reconstruction loss. In the meantime, the output of the encoder, i.e. the d-dimensional feature vector, is submitted to the classification module, which consists of a fully-connected dense layer with softmax nonlinearity and performs the actual classification task. The output of this module, i.e. classification probability, is then compared to the ground-truth label associated with the WSI and a cross-entropy (CE) loss is produced. CE-loss is finally added to the MSE loss to acquire the total loss. By back-propagating on this combined loss function, we train the model to output classification scores that are closer to the tile-level targets while achieving representations of each tile that are independent of image noise (i.e. irrelevant features).

Model hyperparameters and the CNN architecture were determined based on a relevant benchmark analysis from the clinical validation study of a DL model developed for molecular profiling of breast cancer^64^. We adopted the best-performing model’s feature extractor network (based on a “resnet34” architecture^65^) and hyper-parameters to configure a CNN for each biomarker in the current pan-cancer study. Our model selection is further endorsed by a recent benchmarking study comparing weakly-supervised DL approaches for biomarker profiling in computational pathology, where the tile-based DL models are shown to likely outperform newer architectures, such as those based on multi-instance learning, in classification tasks^66^. Each model was trained for 10 epochs using the Adam optimiser with a learning rate of 0.0001. A total of 200 tiles were randomly sampled from each of the training slides and oversampling was applied to the tiles from the underrepresented class to ensure that there is roughly a 50-50 representation of each class during training. During validation, predictions across all tiles were averaged to determine a slide-level prediction. Validation AUC was monitored as the target metric to select the final model during training. Classification scores across images of the same patients were averaged to compute AUC values at patient level.

### Performance characteristics and statistical procedures

Performance of a model was measured with the area under receiver operating characteristic curve (AUC), which plots the relationship between True Positive Rate (TPR) and False Positive Rate (FPR) across different predictive thresholds. An AUC of 0.5 denotes a random model, while a perfect model that can predict all samples correctly yields an AUC of 1. For each biomarker, we reported the performance as the average AUC across the three models (unless otherwise specified), accompanied with the standard deviation (denoted with ± where appropriate).

Statistical significance of results were determined with a two-sided t-test using the “ttest_ind” function from the python scipy-stats library. This is a test for the null hypothesis and assumes that two independent samples have identical expected values and variances. For testing the significance at the group level, the measured AUC values were compared against a randomly-sampled set of values with the same underlying variance. For instance, to measure the statistical significance of the difference between the AUC values acquired for a specific cancer and biomarker type, all AUC values from that group were compared against a set of random AUC values sampled from a distribution with a mean of 0.5 (resembling random performance) and the standard deviation of the group in comparison. To determine the statistical significance of the predictability at biomarker level, we applied the same test on the prediction scores obtained from the true negative and positive cases. Towards this end, classification scores from all three folds of a biomarker were retrospectively combined and the scores from the positive cases and those from the negative class were compared with a two-sided t-test to determine if the difference between the negative and positive predictions were statistically significant. The resulting p-values were corrected for multiple testing using the Benjamini-Hochberg procedure with a false discovery rate (FDR) of 0.05. All biomarkers with an adjusted p-value of less than 0.05 were considered statistically significant.

The experiments and analyses in this work were carried out with python 3.6 and the following python libraries: torch (1.9.0), torchvision (0.10.0), google-cloud-storage (1.32.0), openslide-python (1.1.1), pillow (6.0.0), opencv (3.4.2), tensorboard (1.15.0), numpy (1.18.1), pandas (1.3.5), seaborn (0.11.0), scipy (1.4.1), scikit-learn (0.22.1), statsmodels (0.11.0), and matplotlib (3.5.1). For the impact of variables on the predictive performance analysis (**Extended Data Figure 7**), we used scikit-learn (0.22.1)’s random forest regressor (RFR) implementation with default parameters.

## Acknowledgement

The results shown in this study are partially based upon data generated by the TCGA Research Network (https://www.cancer.gov/tcga) the Clinical Proteomic Tumour Analysis Consortium (NCI/NIH, https://proteomics.cancer.gov/programs/cptac). The authors would also like to thank Oscar Maiques for the discussion prior, during and after the writing of this publication.

## Data availability

TCGA whole slide images are available at https://portal.gdc.cancer.gov/ . Genetic, transcriptomic, proteomic, and clinical data used to generate biomarker profiles for cases in the TCGA cohorts are available at https://portal.gdc.cancer.gov/ and https://cbioportal.org/. Clinically relevant driver genes are available at https://cancervariants.org/. Personalised genome-scale metabolic models used for metabolic pathways are available at https://www.ebi.ac.uk/biomodels/pdgsmm/index/. CPTAC whole slide images are available at https://wiki.cancerimagingarchive.net/display/Public/CPTAC+Imaging+Proteomics/. Genetic data used to generate biomarker profiles for cases in the CPTAC cohorts are available at https://portal.gdc.cancer.gov/.

## Authors’ contribution

P.R-L. conceptualised and led the research. P.R-L. and S.A. designed and performed the study experiments. S.A., X.L., J.H. and C.B. developed the methods validated in the experiments. J.N.K, D.M. and S.S. provided clinical inputs for the research. S.A., J.S., A.G., J.H. implemented the software needed to run the experiments. P.R-L., S.A., D.M. and S.S. analysed the experimental results. S.A., P.R-L., D.M. and S.S. wrote the manuscript. Everyone reviewed the manuscript.

## Extended Data

**Extended Data Figure 1.**
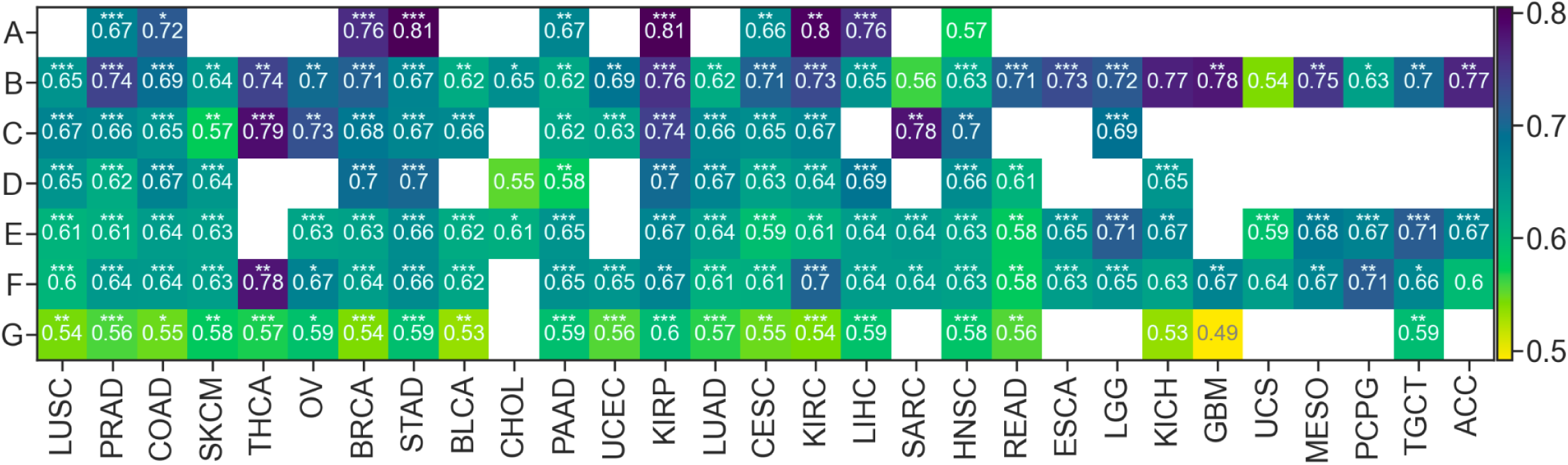
| Predictability performance for each malignancy across all biomarker types. Average performance (AUC) across all cancer studies confounded by biomarker type are shown in a heatmap. Empty cells correspond to having no data for those cancer-biomarker groups. Asterisks within heatmap cells indicate the statistical significance of the performance difference between AUC values of a subgroup against randomly sampled values of the same underlying distribution (no asterisk: not significant (n.s.), *: p < 0.05, **: p < 0.01, ***: p < 1e-05). DLBC, UVM, and THYM were excluded from the analysis due to only constituting one to seven valid targets across all biomarker types. The following coding was used to abbreviate the biomarker types: **A** for standard clinical biomarkers; **B** for clinical outcomes and treatment responses; **C** for under-/over-expression of proteins; **D** for gene signatures and molecular subtypes; **E** for under-/over-expression of driver genes; **F** for driver SNV mutations; and **G** for metabolic pathways. Overall, we obtained a better-than-random average performance across all biomarker types (i.e. mean AUC > 0.5), except for metabolic pathways in glioblastoma multiforme (GBM). Among the cancer types targeting standard clinical features **(A)**, kidney renal papillary cell carcinoma (KIRP, AUC: 0.805 ±0.132, p < 0.01) and stomach cancer (STAD, AUC: 0.805 ±0.084, p < 1e-05) had the top average performance, followed by clear cell renal cell carcinoma (KIRC), breast adenocarcinoma (BRCA), and colon cancer (COAD), each with a mean AUC over 0.7. Multiple cancer types showed a relatively good performance with average AUCs above 0.7 especially when considering the predictability of the clinical outcomes and treatment responses **(B)**, where the performances of such predictions were among the highest across studies. Among them, the most notable studies were kidney renal papillary cell carcinoma (KIRP), adrenocortical carcinoma (ACC), glioblastoma multiforme (GBM), and kidney chromophobe (KICH) with average AUCs reaching as high as 0.777. For genomic, transcriptomic, and proteomic biomarkers **(C, E, F)**, the performances within individual cancer types were primarily consistent with their corresponding general trend. The highest performances were observed in thyroid carcinoma (THCA) and sarcoma (SARC) for the prediction of proteomic expression status with average AUCs around 0.78; in lower-grade glioma (LGG) and testicular germ cell tumours (TGCT) for the predictability of transcriptomic biomarkers with AUCs slightly above 0.7; and in kidney renal clear cell carcinoma (KIRC), pheochromocytoma/paraganglioma (PCPG), and thyroid carcinoma (THCA) for the detection of genetic alterations in driver genes with average AUCs ranging from 0.705 to 0.779. Top-performing cancers from the gene signatures and molecular subtypes **(D)** were breast cancer (BRCA), gastric cancer (STAD), and kidney renal papillary cell carcinoma (KIRP), each having an average AUC of 0.7. Cancer-specific performance for the prediction of metabolic pathways **(G)** was on par with its overall low-performance, with none of them yielding an average AUC above 0.6.

**Extended Data Figure 2.**
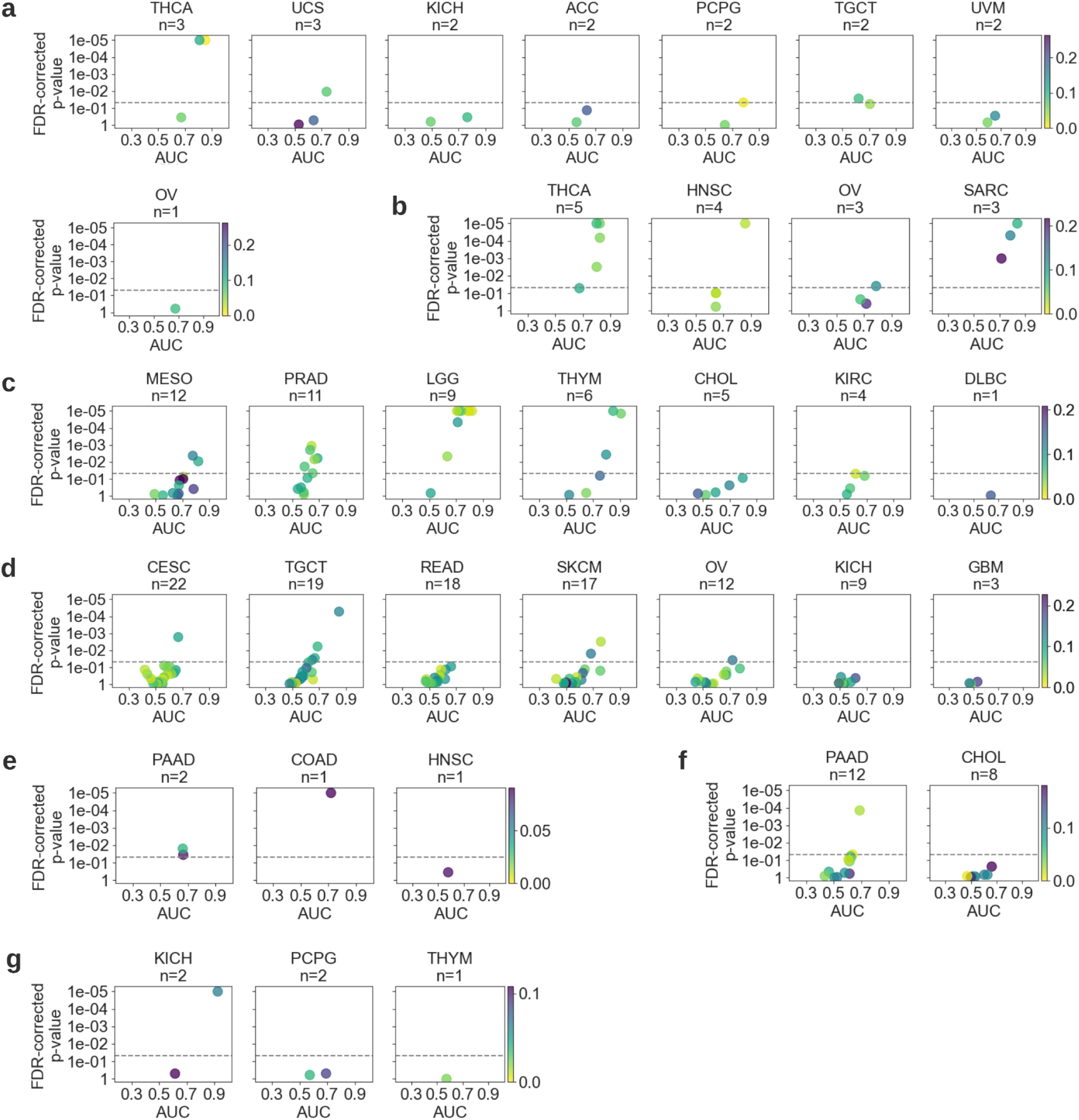
| Performance for biomarker predictability across selected cancer types. Scatter plots showing the performance of each model trained to predict biomarkers across different categories, namely, **(a)** driver SNV mutations, **(b)** protein under-/over-expression status, **(c)** under-/over-expression of driver genes at transcript level, **(d)** metabolic pathways, **(e)** standard clinical biomarkers, **(f)** gene signatures and subtypes, and finally, **(g)** clinical outcomes and treatment responses. Only the cancer types that were not included in the main figures due to space limitations are shown here. Please refer to the caption of Figure 3 for a detailed explanation of the visualisation.

**Extended Data Figure 3.**
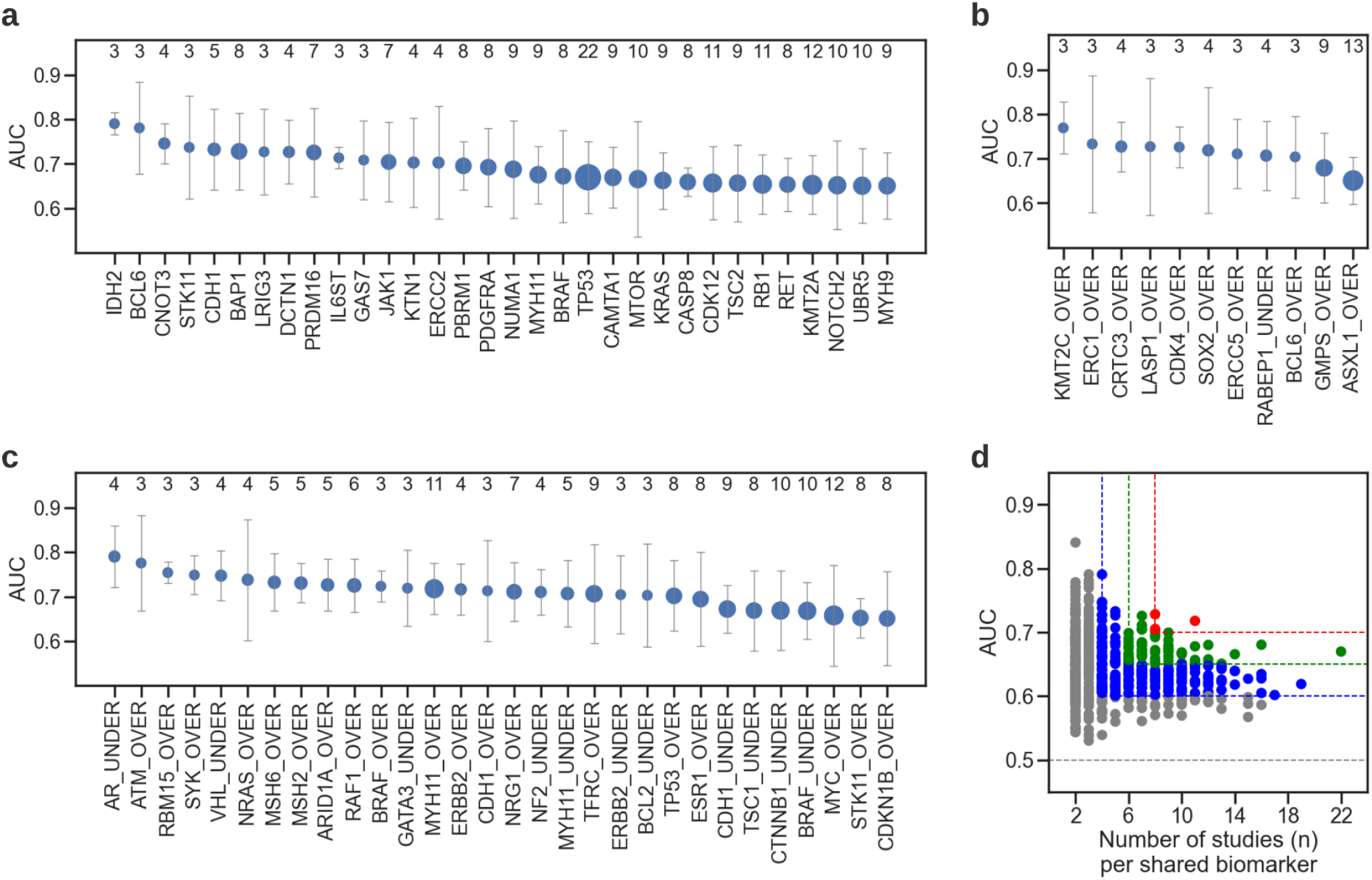
| Biomarkers that were predictable in multiple cancers. **(a-c)**: Driver mutations, transcriptome and protein expression status can be predicted in at least three and seven cancer types with an average AUC of 0.7 or 0.65, respectively. Whiskers show the standard deviation of AUC across cancers. The size of a marker represents the frequency of a biomarker. The exact number of appearances (n) of a biomarker across multiple cancers is further shown in the secondary x-axis. **(a)** Alterations in *TP53* were detectable in almost all cancer types, with 7 of them having an AUC of at least 0.7 and 14 of them showing AUCs greater than 0.65. Other genes with a cross-cancer AUC of at least 0.7 were *BAP1* (mutations predictable in eight cancers), *PRDM16* and *JAK1* (mutations predictable seven cancers), and *CDH1* (mutation predictable five cancers). *CDK12, RB1, MTOR, NOTCH2, UBR5, and KMT2A*, are also worth mentioning with their mutations being detectable in at least 10 different cancers with a mean AUC of 0.65. **(b)** Over-expression of *KMT2C* had a consistently high prediction rate with AUCs ranging 0.733-0.837 in kidney renal papillary cell carcinoma, ovarian serous cystadenocarcinoma, and testis cancer. Other notable genes that were detectable across multiple cancer types at over-expression levels were *ERC1, CRTC3, LASP1, CDK4, SOX2, ERCC5, BCL6* and the under-expression status of *RABEP1*, each being predicted in three or more different cancers. **(c)** Over-expression of *MYH11* was detected in 7 out of 11 cancer types with AUCs ranging from 0.705 to 0.809. Other notable genes associated with protein over-expression were *TFRC, TP53, MSH2, MSH6, ARID1A, RAF1*, and NRG1, showing a high-predictability in at least five malignancies, with AUCs reaching 0.942. Under-expression of proteins encoded by *MYH1, NF2, VHL*, and *AR* were also predictable in at least four cancers, with AUCs reaching 0.856. **(d)** Scatter plot showing the AUC values of genetic alterations, as well as transcriptome and protein expression status that can be predicted in multiple cancer types. Areas outlined with red, blue, and green lines mark the zones with high predictability and frequency of appearance. The red zone corresponds to the biomarkers with an AUC of at least 0.7 and a frequency of eight and above. The green zone corresponds to the biomarkers with an AUC of at least 0.65 and a frequency of six and above. For blue, AUC and frequency are limited to 0.6 and four, respectively.

**Extended Data Figure 4.**
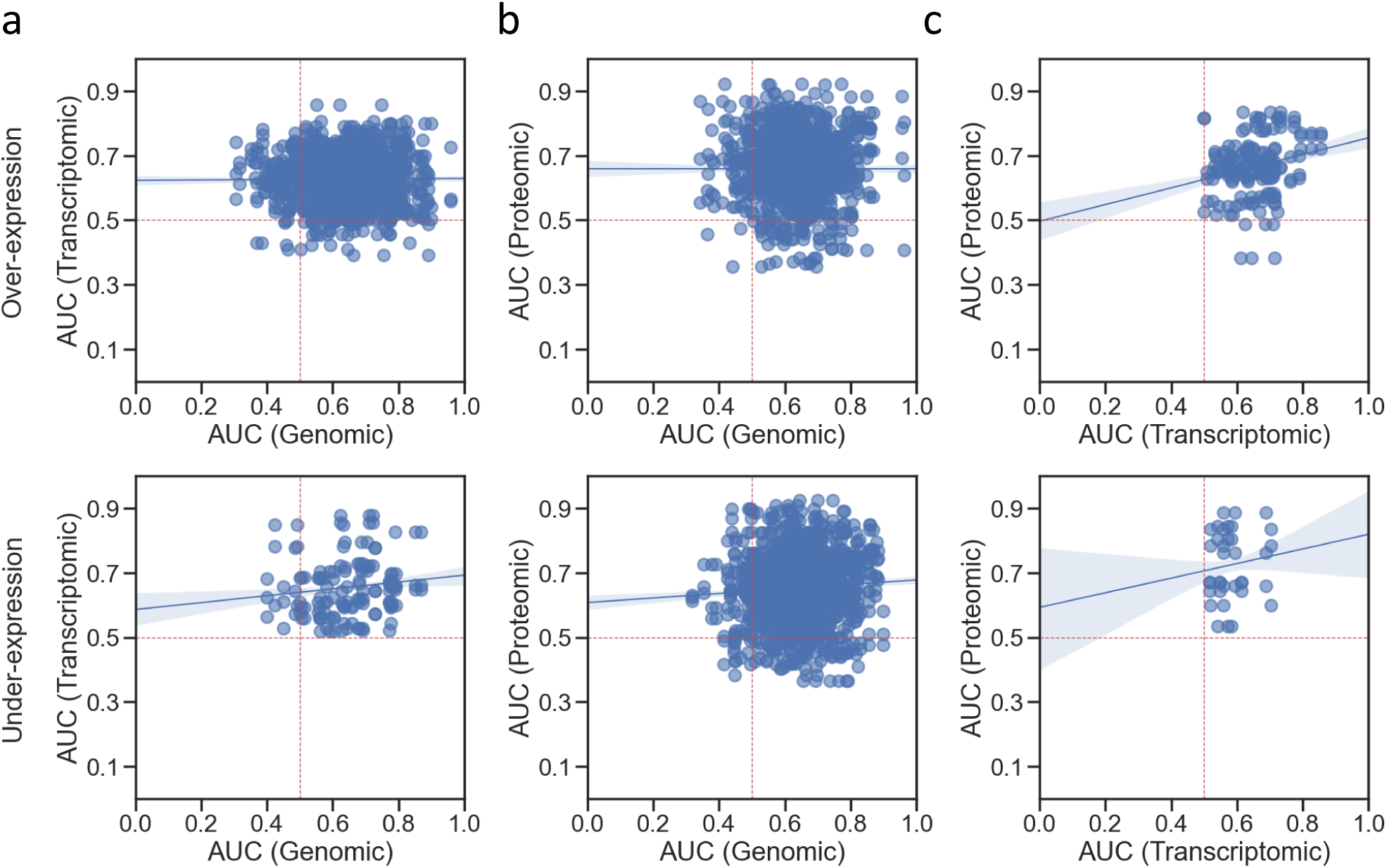
| Cross-correlation of the detectability of molecular alterations across genomic, transcriptomic and proteomic biomarkers. For this analysis, driver genes with a valid genomic, proteomic, and transcriptomic profile (**Online Methods: Biomarker acquisition**) across all cancer types were identified and cross-correlated with Pearson correlation coefficient. Since there existed three models per biomarker, each model in a given omic type was compared to all the three models in the other omic type, yielding nine comparisons per gene. **(a-b)** Both at the transcriptomic and proteomic expression levels there was no correlation between the predictability of genetic alterations and under- or over-expression status associated with them, except for under-expressed transcriptomes and genetic alterations (Pearson correlation: 0.131, p<0.01). **(c)** We measured a positive correlation of 0.227 (p<1e-05) and 0.124 (p=0.203) between the predictability of genes associated with transcriptomes and proteins at over- (*top*) and under- (*bottom*) expression levels, respectively.

**Extended Data Figure 5.**
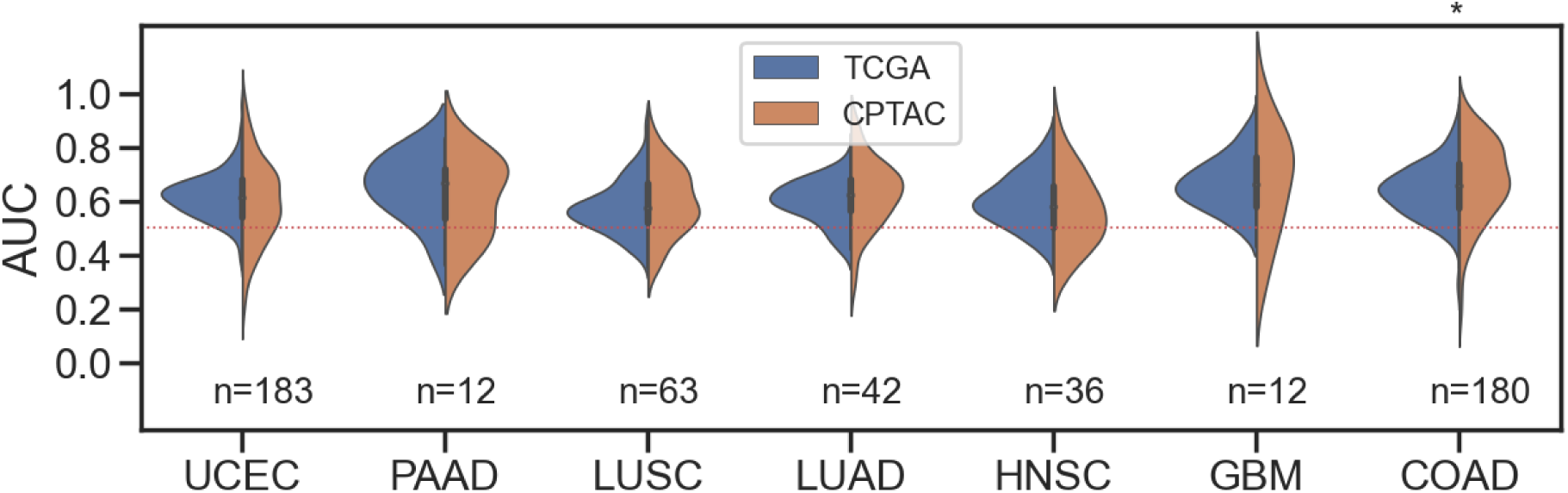
| Reproducibility of pan-cancer predictability on external dataset. We repeated our experiments using CPTAC data, to show that a comparable performance can be achieved on a different dataset. The data only contains driver SNV mutations due to both TCGA and CPTAC relying on the same set of driver genes, hence exhibiting a relatively large overlap. A total of 176 driver genes (corresponding to 528 models) across seven cancer types had qualified mutation data in both datasets (**Online Methods: Biomarker acquisition)**. Number of models validated per cancer type is shown under each violin plot for each cohort. The investigated cancers were uterine corpus endometrial carcinoma (UCEC), pancreatic ductal adenocarcinoma (PAAD), lung squamous cell carcinoma (LUSC), lung adenocarcinoma (LUAD), head and neck cancer (HNSC), glioblastoma multiforme (GBM), and colon adenocarcinoma (COAD). Asterisk atop a violin plot indicates the difference between AUC values of two chorts being statistically significant (i.e. p < 0.05). Here, all biomarkers but those from COAD had comparable AUC distributions in TCGA and CPTAC cohorts.

**Extended Data Figure 6.**
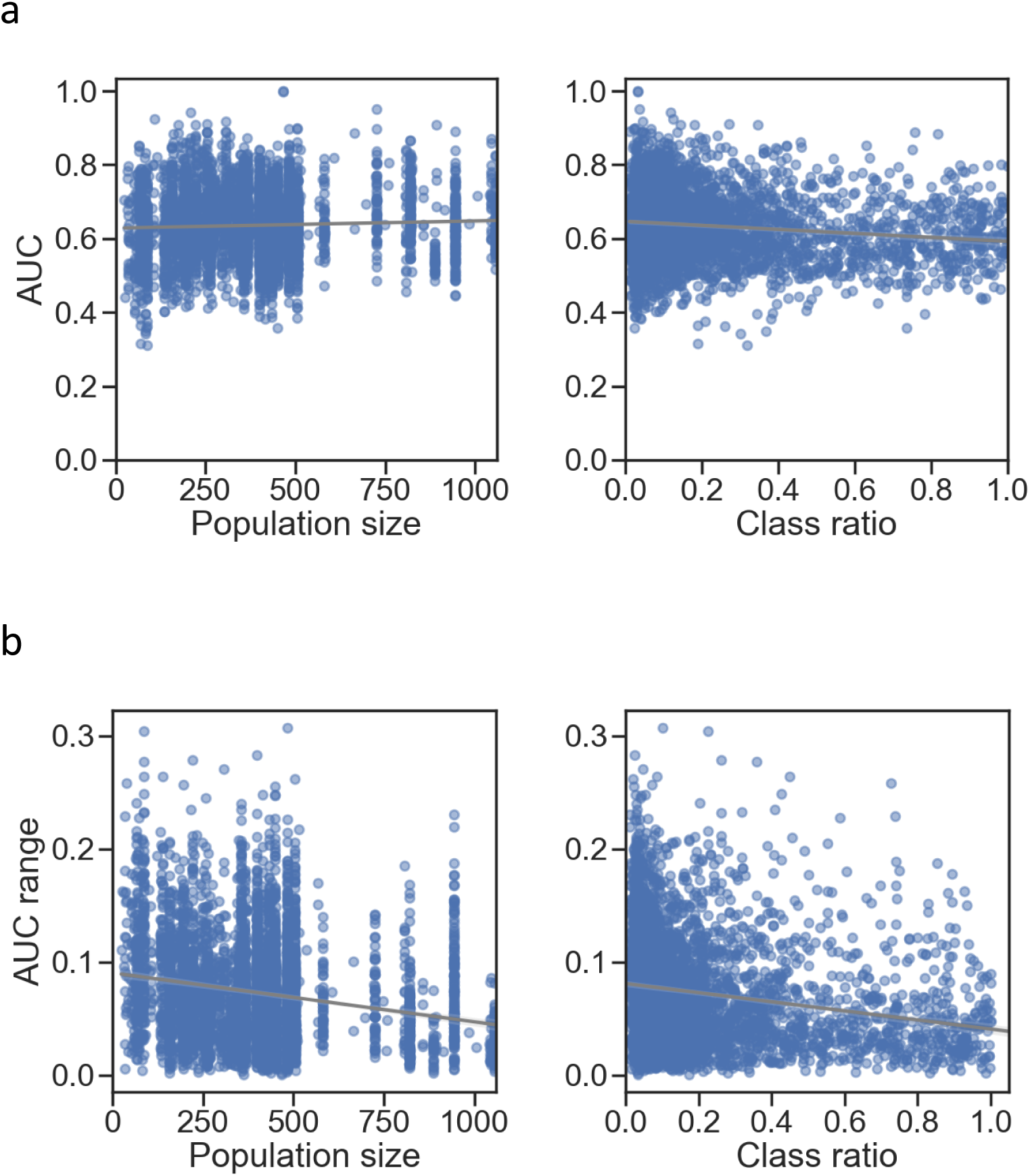
| Impact of sample size and positive class ratio on the prediction performance. We evaluated the potential influence of the number of samples and the class proportions on the prediction performance, by correlating population sizes and class ratios with **(a)** the biomarker AUC values and **(b)** the variability in biomarker performance as measured by standard deviation across the AUCs of CV folds. Population size corresponds to the number of total samples per biomarker and class ratio is computed as the size of underrepresented class over the size of other class (i.e. a ratio close to 1 denotes a perfectly balanced class distribution, whereas a ratio close to 0 means a severely unbalanced dataset). Considering the amount of steps involved in biomarker acquisition for each omic type (**Extended Data: Biomarker acquisition**) and the diverse number of diagnostic slides available for each cancer (**Extended Data Table 2**), the population size and class distributions per biomarker varied quite significantly across different malignancies (**Extended Data Figure 8**). **(a)** Pearson correlation coefficient (PCC) between the sample size and the AUC values was 0.046 (p-value < 0.01), indicating no linear relationship between the two variables. Similarly, a PCC of -0.139 (p-value < 1e-05) was obtained for the class ratio, showing almost no impact of class on predictiability. **(b)** A weak negative relationship was observed for the AUC variability when it was correlated with population size (PCC: -0.196, p < 1e-05) and class ratios (PCC: -0.197, p < 1e-05). This might indicate that the performance tends to become more stable with an increasing number of samples and a more balanced dataset.

**Extended Data Figure 7.**
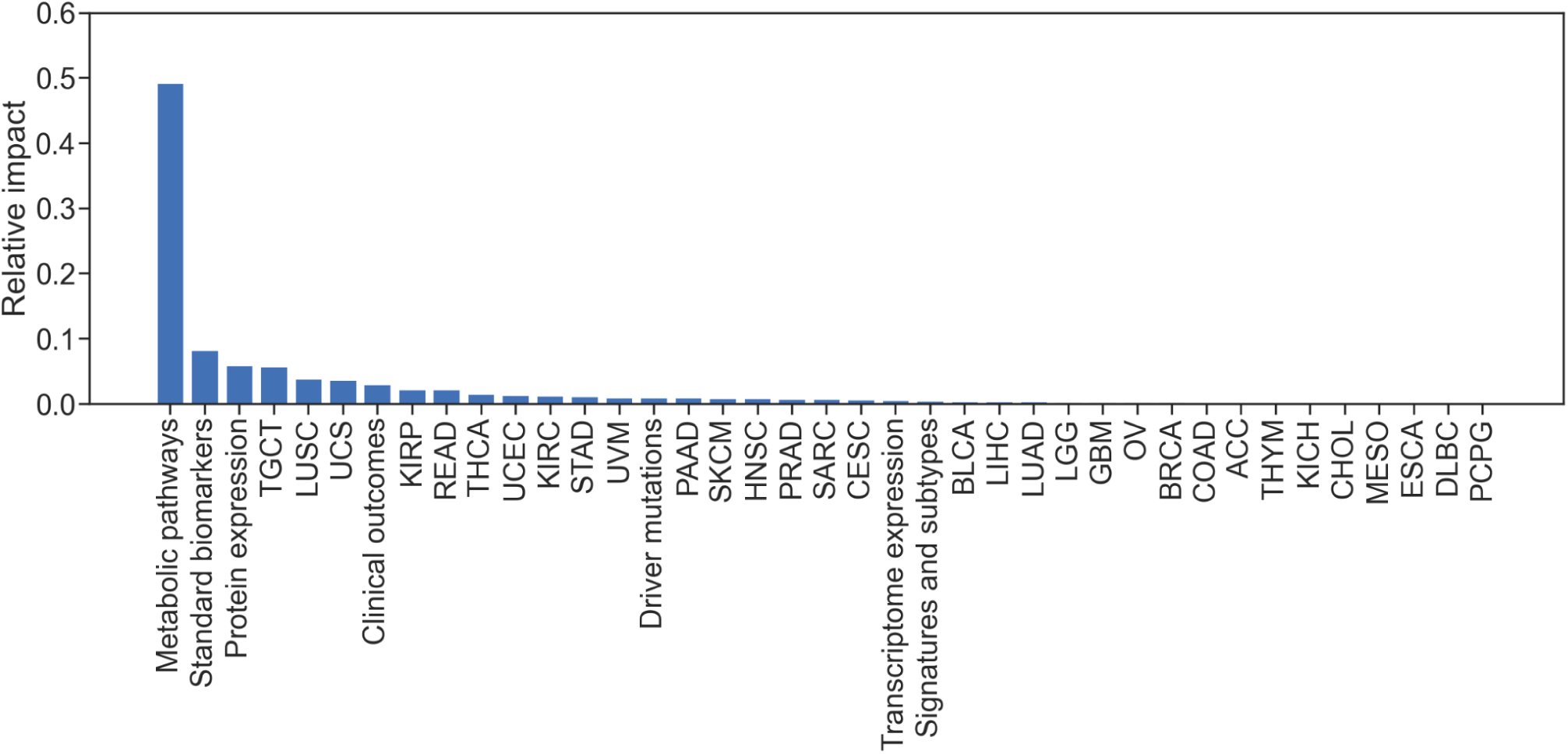
| Impact of other factors on the predictive performance. Considering the very low impact of class ratio and population size on the biomarker performance (**Extended Data Figure 8**) we performed a simple experiment to assess the importance of other factors such as the type of cancer and biomarker class for predictability. These categorical variables were converted into variable indicator variables via one hot encoding to obtain a 39-dimensional feature matrix. A random forest regression (RFR) model was fitted on this data to predict the AUC values. During training, the RFR model assigns a weight to each feature, which, in turn, can be used to estimate the “predictiveness” of a variable for regressing the AUC values). Overall, the combined impact of the omic types was the largest, with metabolic pathways being the most important feature for predictive performance. The impact of cancer types, on the other hand, was rather limited.

**Extended Data Figure 8.**
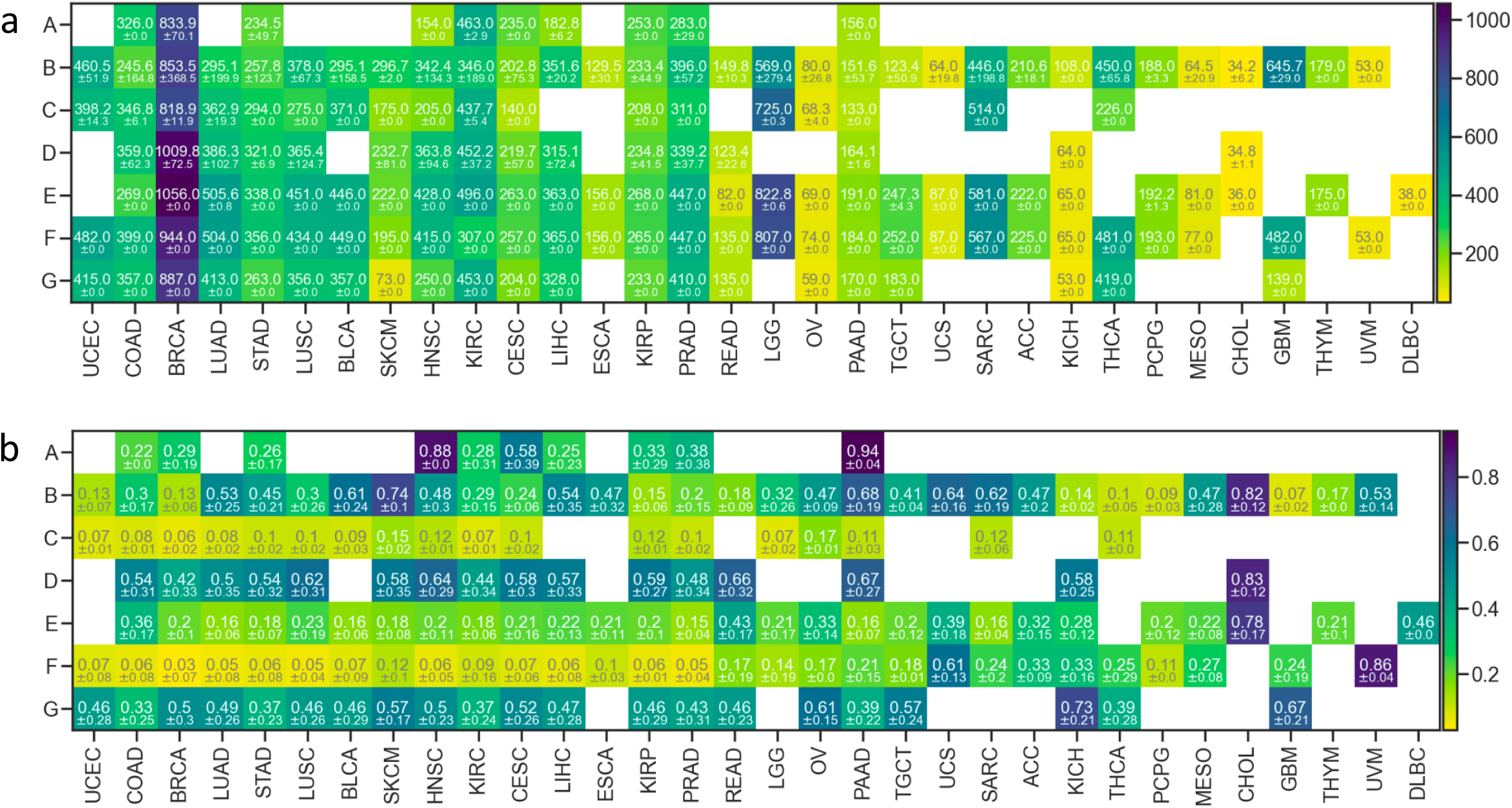
| Average population size and class ratio across all cancer and biomarker types. **(a)** Average population size and **(b)** average class ratio across all cancer and biomarker types are shown in a heatmap. Population size corresponds to the number of total samples per biomarker and class ratio is computed as the size of underrepresented class over the size of the other class (i.e. a ratio close to 1 denotes a perfectly balanced class distribution, whereas a ratio close to 0 means a severely unbalanced dataset). Empty cells indicate no data for those cancer-biomarker groups. The following coding was used to abbreviate the biomarker types: **A** for standard clinical biomarkers; **B** for clinical outcomes and treatment responses; **C** for under-/over-expression of proteins; **D** for gene signatures and molecular subtypes; **E** for under-/over-expression of driver genes; **F** for driver SNV mutations; and **G** for metabolic pathways.

**Extended Data Figure 9.**
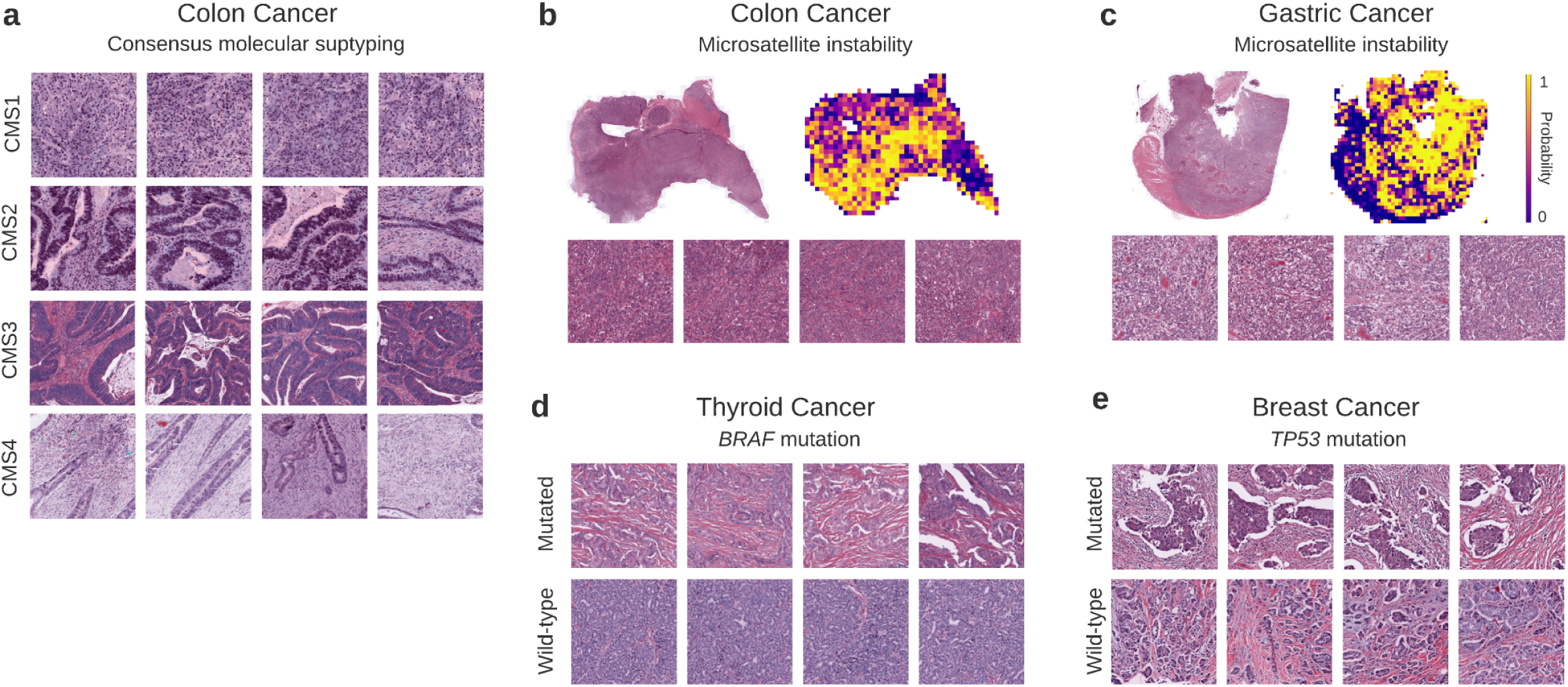
| Highest scoring tiles from certain biomarkers in colon, gastric, thyroid, and breast cancers. **(a)** The top-ranking tiles from the consensus molecular subtypes (CMS) of colon cancer (i.e. CMS1, CMS2, CMS3, and CMS4) show distinct morphological features. One can see lymphocytic infiltration patterns in CMS1, well-differentiated glandular structures for CMS2-3, and high stromal content in CMS4 tiles. **(b-c)** Highly predicted tiles from colon and gastric cancer patients showing morphological traits associated with microsatellite instability. **(d-e)**. The-highest ranking tiles for the prediction of *BRAF* mutation in thyroid carcinoma **(d)** and *TP53* mutation in breast cancer **(e)** compared to their wild-type counterparts.

**Extended Data Figure 10.**
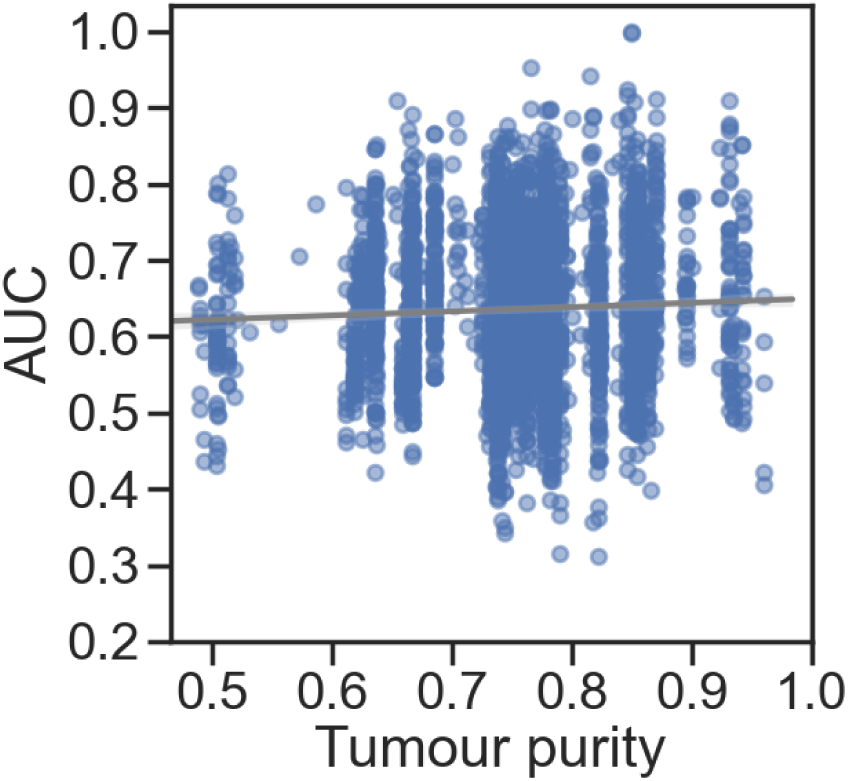
| Impact of tumour purity on the prediction performance. We evaluated the potential impact of the tumour composition on the performance of the predictive models, by correlating tumour purity with the biomarker AUC values. Tumour content was approximated by the percentage of tumour cells in whole slide images and averaged across samples for each biomarker. Scatter plot shows how the average tumour purity is correlated with AUC for all biomarkers evaluated in this study. Pearson correlation coefficient (PCC) between tumour purity and the AUC values was 0.044 (p-value < 0.01), showing no apparent relationship between the two variables. This might indicate that tumour purity is unlikely to bias the predictability from H&E images.

**Extended Data Table 1.**
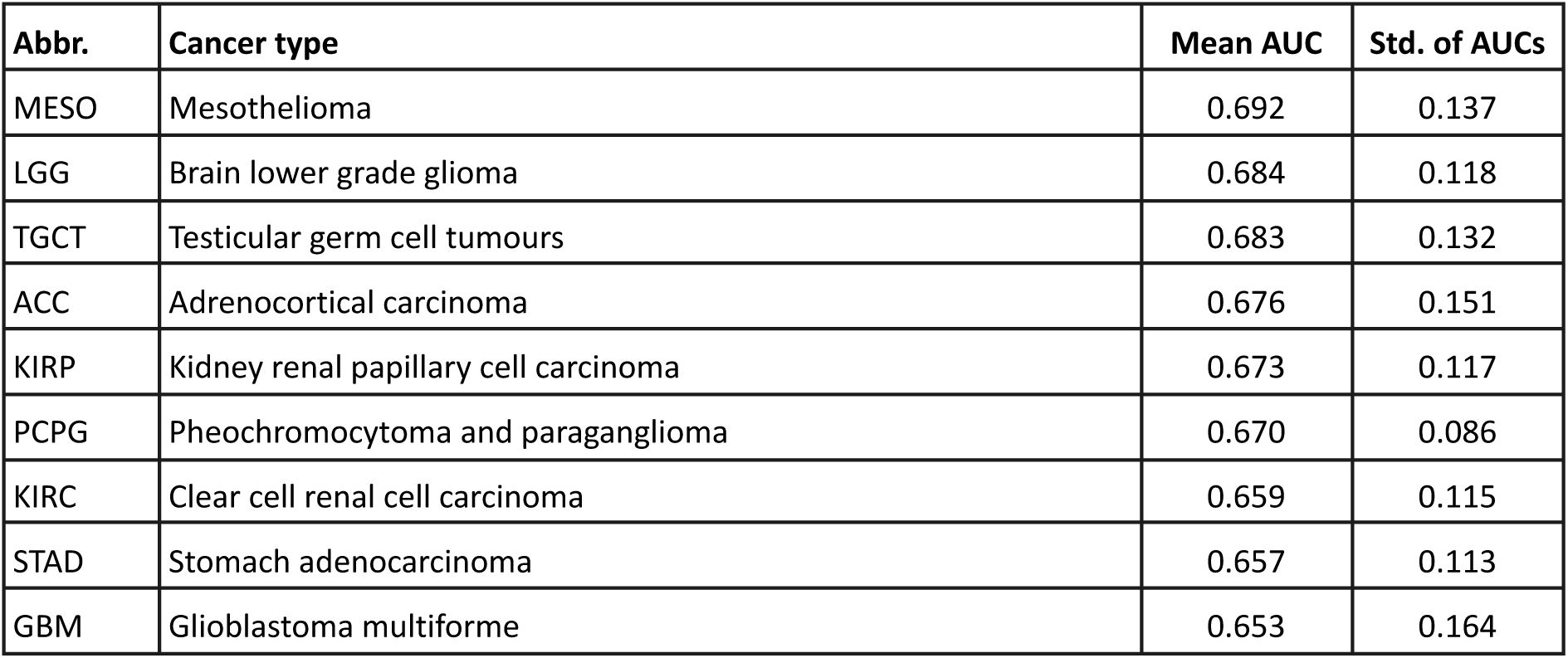
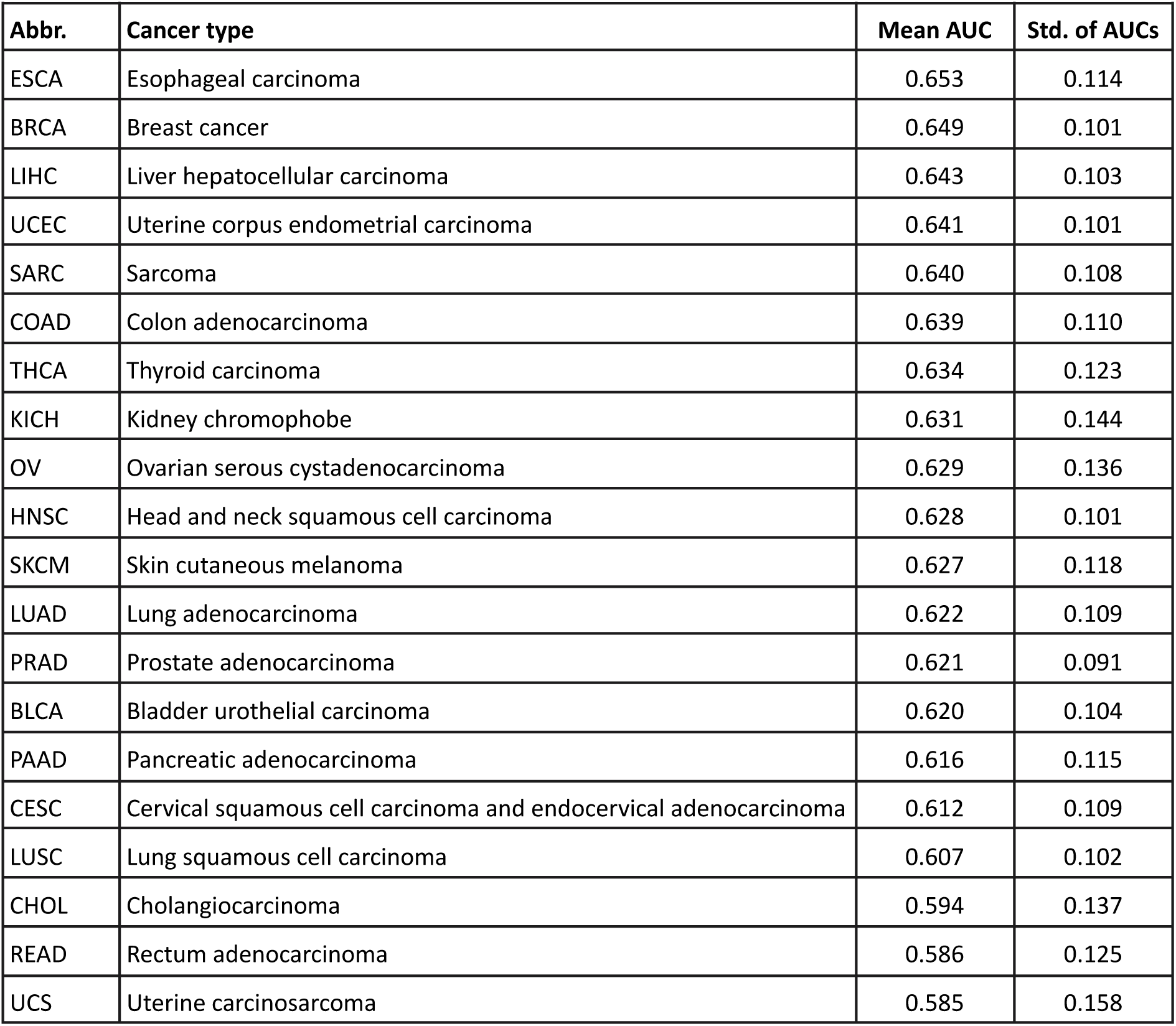
| Average performance and standard deviation for all cancer types. DLBC, UVM, and THYM were excluded from the table due to only constituting one to seven valid targets across all biomarker types.

**Extended Data Table 2.**
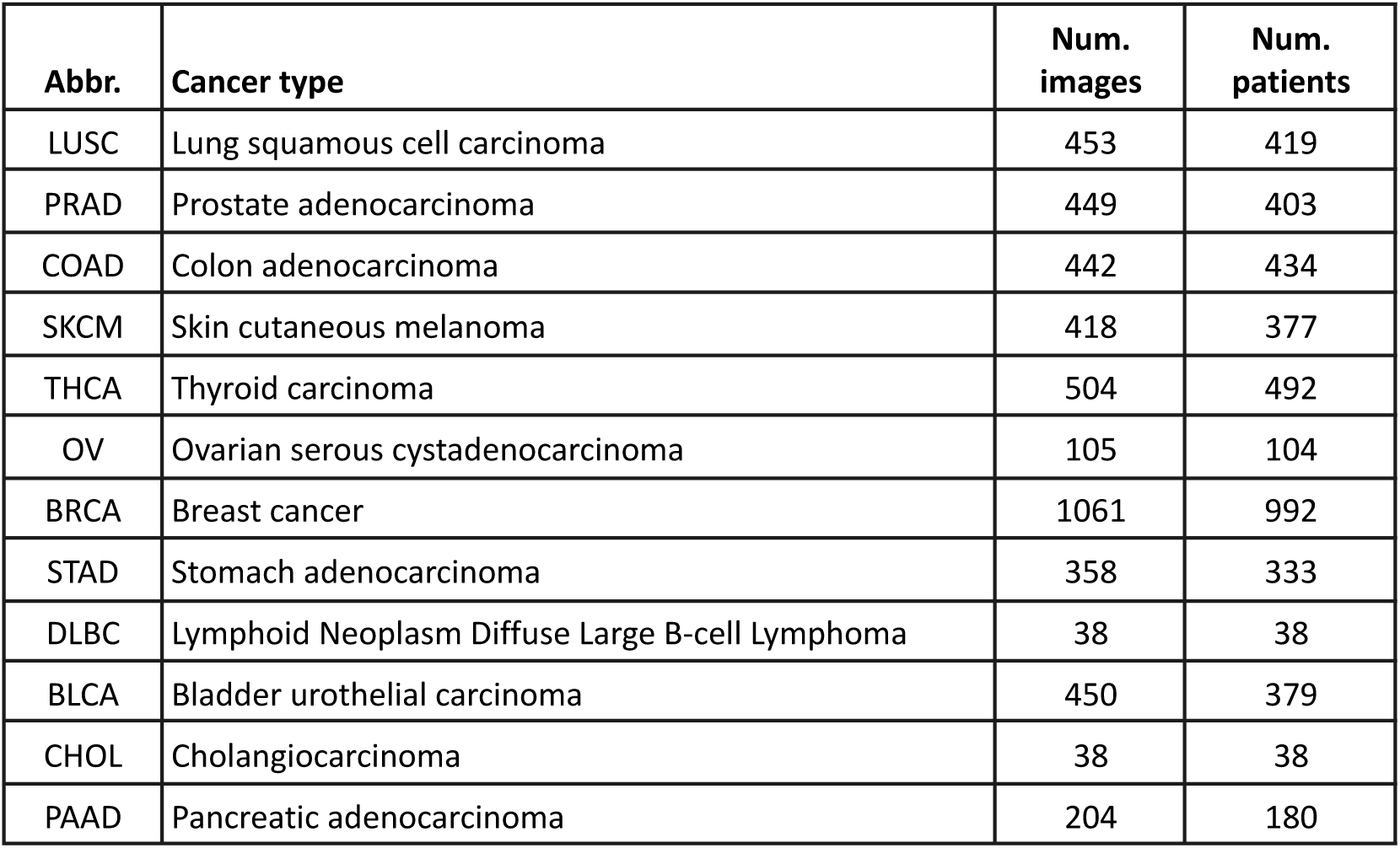
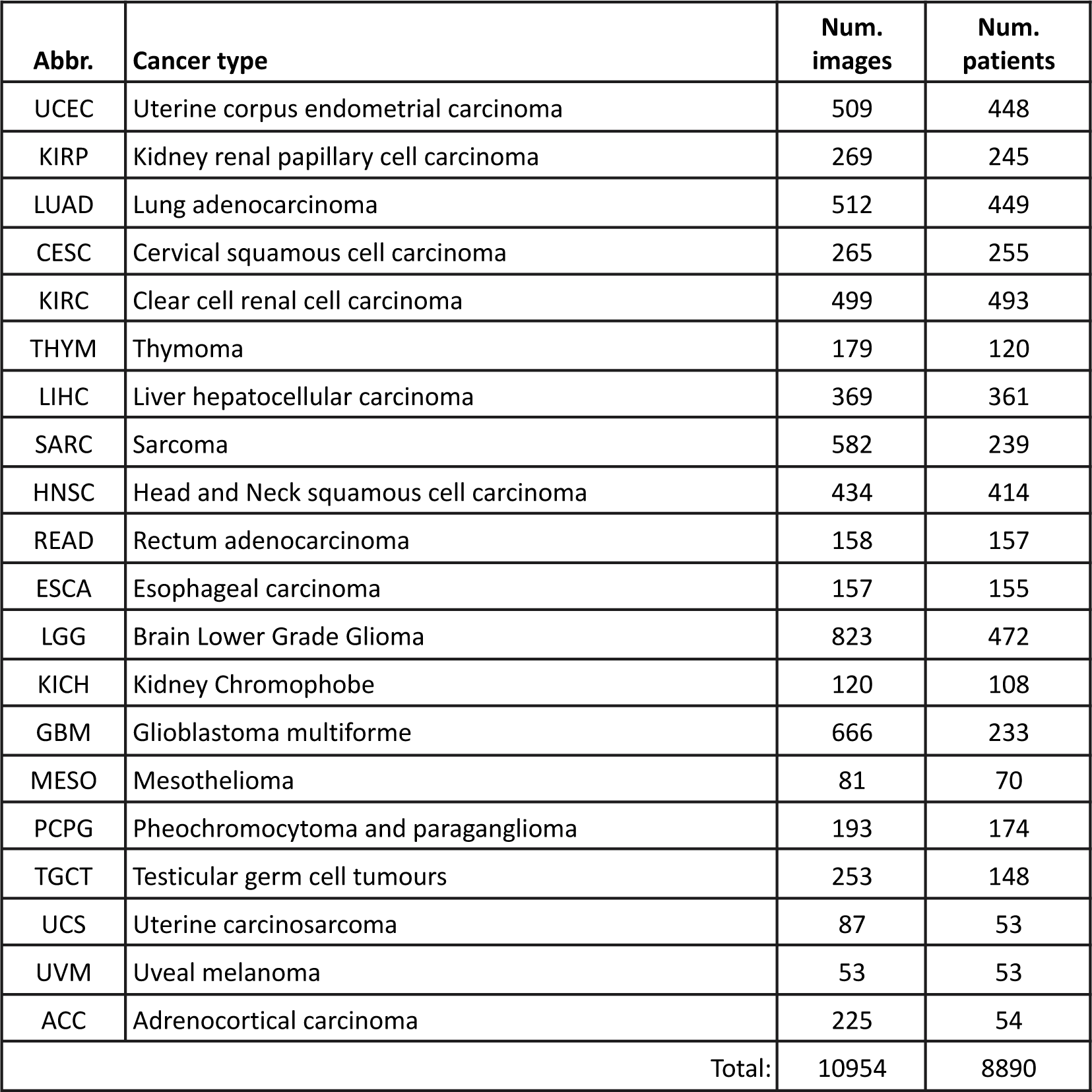
| Number of images and patients included in the study across all cancer types. Cancer names corresponding to abbreviations are provided in Extended Data Table 1.

**Extended Data Table 3.** | Number of images and patients included in the CPTAC experiment. Only the cancers that have comparable biomarkers in the TCGA dataset were considered. The abbreviations used in the CPTAC and TCGA resources for the same cancer type are provided in the first two columns, respectively.

| **Abbr.** | **TCGA-like abbr.** | **Cancer type** | **Num. images** | **Num. patients** |
| --- | --- | --- | --- | --- |
| UCEC | UCEC | Uterine corpus endometrial carcinoma | 591 | 247 |
| PDA | PAAD | Pancreatic ductal adenocarcinoma | 382 | 168 |
| LSCC | LUSC | Lung squamous cell carcinoma | 689 | 211 |
| LUAD | LUAD | Lung adenocarcinoma | 669 | 224 |
| HNSCC | HNSC | Head-and-neck cancer | 268 | 112 |
| GBM | GBM | Glioblastoma multiforme | 510 | 189 |
| COAD | COAD | Colon adenocarcinoma | 372 | 178 |
| Total: |  |  | 3481 | 1329 |

